# T-cell Receptor (TCR) Targeting with Multivalent T-cell Engagers

**DOI:** 10.64898/2026.05.17.725710

**Authors:** Debasmita Paul, Abhishek Kulkarni, Freddys Rodriguez, Declan G Dahlberg, Lakmal Rozumalski, Carston R Wagner

## Abstract

T-cell engagers (TCEs) for cancer immunotherapy have traditionally relied on high affinity single chain fragment variable (scFv) domains to target CD3, specifically the ε chain, for the activation of T-cells. Despite their clinical success, there have been reports of TCEs driving systemic toxicity, non-specific T-cell activation, on-target off-tumor effects, and severe inflammation due to cytokine release. To address these limitations, we designed multivalent TCEs using Chemically Self-Assembled Nanorings (CSANs) that target the α/β constant region of the T-cell receptor (TCR) in the TCR/CD3 complex using a moderate affinity αTCR nanobody (αTCR_VHH_). Nanobodies offer superior physical and chemical properties over scFvs-including higher solubility, stability and lower production cost-making them increasingly popular as structural units of TCEs. We compared the efficacy and safety profile of this moderate affinity, nanobody-based TCR binder against high affinity αCD3_scFv_ based CSANs across EGFR and PSMA expressing solid tumor models. While the αCD3_scFv_ CSANs offered potent cytotoxicity, they also induced antigen independent T-cell activation bypassing the requirement of tumor crosslinking for cytotoxicity. In contrast the αTCR_VHH_ CSANs required strict antigen engagement to trigger cytotoxicity, significantly reducing non-specific T-cell activation and thus enhancing the safety profile. Although the initiation of cytotoxicity was kinetically slower than the αCD3_scFv_ counterpart, αTCR_VHH_ CSANs achieved comparable end point cytotoxicity across multiple antigen densities, as well as in 3D tumor spheroids. Through this study we demonstrate the applicability of nanobodies as T-cell targeting domains, enhanced specificity and safety of moderate affinity T-cells binders and the diversification of T-cell targeting epitopes without compromising the efficacy of TCEs.

**Abstract Graphic:** 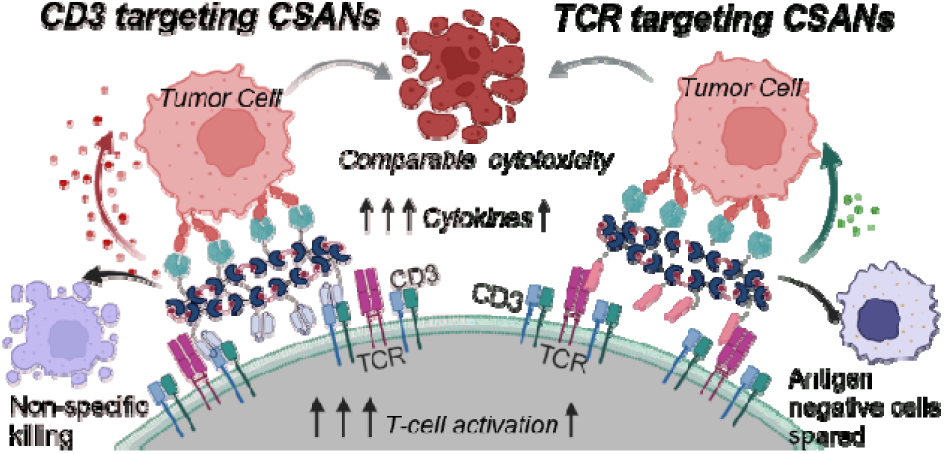

## 1. INTRODUCTION

T-cell engagers (TCEs) have garnered increasing prominence in recent years as a transformative ‘off-the-shelf’ platform in immunotherapy, where the patient’s immune system is harnessed to target and eliminate cancer cells. Unlike checkpoint inhibitors that release the ‘brakes’ on the immune system, activating tumor specific T-cells or CAR-T therapies that require complex genetic engineering of the patient’s immune cells, TCEs act directly to activate anti-tumor T-cell responses. TCEs are typically fusion proteins engineered to have at least two binding domains: one specific for a Tumor Associated Antigen (TAA) on the cancer cell surface, and the other specific for the TCR/CD3 complex, specifically for the CD3 subunit, on the T-cell surface^1^. By physically bridging the T-cells to tumor cells, TCEs force the formation of an immunological synapse. This proximity triggers the downstream signaling cascades of the T-cell, resulting in T-cell activation, release of cytotoxic molecules such as perforins and granzymes, and the subsequent lysis of the tumor cells.

A critical advantage of this platform is the ability to bypass the Major Histocompatibility Complex (MHC) restriction. Many tumors adopt mechanisms of immune evasion by downregulating the MHC expression or impairing antigen processing and presentation, thus effectively being invisible to immune surveillance^2^. By circumventing this requirement, TCEs can enable activation of the T-cells against tumor cells regardless of the MHC status.

The first FDA approved TCE, Blinatumomab, which is used to treat B-cell precursor acute lymphoblastic leukemia (B-ALL), set the stage for the current development of TCEs for hematological malignancies.^3–6^ Nevertheless, the translation of TCE efficacy to solid tumors has been challenging. Conventional high-affinity TCEs often fail to discriminate between the high antigen expression on tumors compared to the typically lower expression on normal tissues, leading to on-target off-tumor toxicity, and potentially life-threatening cytokine release syndrome (CRS).^7–9^ Further, solid malignancies present a hostile and immunosuppressive microenvironment that is populated by Regulatory T-cells (T_regs_), myeloid-derived suppressor cells (MDSCs), and anti-inflammatory cytokines that dampen the proper functioning of the pro-inflammatory immune cells. The physical barrier posed by the complex stroma often prevents a sufficient number of T-cells from reaching the tumor core.^10^ Structural modifications to TCEs that facilitate better penetration into the dense and solid tumor microenvironment have often negatively impacted their stability and half-life.^1, 4, 11^ Thus, the development of solid tumor TCEs depends on tuning T-cell activation to enhance specificity and tissue penetration by attenuating maximal potency and affinity.

Currently, the majority of the TCEs approved or under development rely on single-chain fragment variable (scFv) engagement of the CD3 subunit of the TCR/CD3 complex. While scFvs retain the specificity of the parental antibody, they generally have reduced biophysical stability that can lead to aggregation hindering large scale manufacturing.^12^ ^13, 14^ ^15, 16^ To address these limitations, single domain antibodies-specifically Variable Heavy Chain antibodies (V_HH_) or nanobodies offer a compelling alternative. Nanobodies are the smallest known antigen-binding domain to date and hence offer excellent penetration in solid tissues, making them candidates for diagnostic and therapeutic applications. In addition, they have improved structural and biophysical stability than scFvs and thus are less prone to aggregation and more easily prepared by bacterial expression.^17–21^

In this study, we leveraged our chemically self-assembled nanoring (CSANs) platform to engineer a multi-valent bispecific T-cell engager that integrates the structural advantages of nanobodies with tunable T-cell activation (Figure 1). Each monomer in the multimeric CSAN is composed of two dihydrofolate-reductase fusion proteins DHFR^2^ linked by a single glycine linker.^22^ Each DHFR^2^ unit is fused to binding ligand that either bind to T-cells or TAAs. Upon addition of the chemical dimerizer, bis-MTX, the DHFR^2^ fusion proteins spontaneously self-assemble to octameric nanorings, generating multi-valent CSANs. This multivalent architecture allows for the generation of multivalent bispecific TCE.^23^ By varying the ratio of the T-cell binding domains to TAA binding domains the signal strength and resultant potency of the CSANs TCE can be varied that have shown potent *in vitro* and *in vivo* T-cell targeted anti-tumor activity.^24, 25^

**Figure 1.**
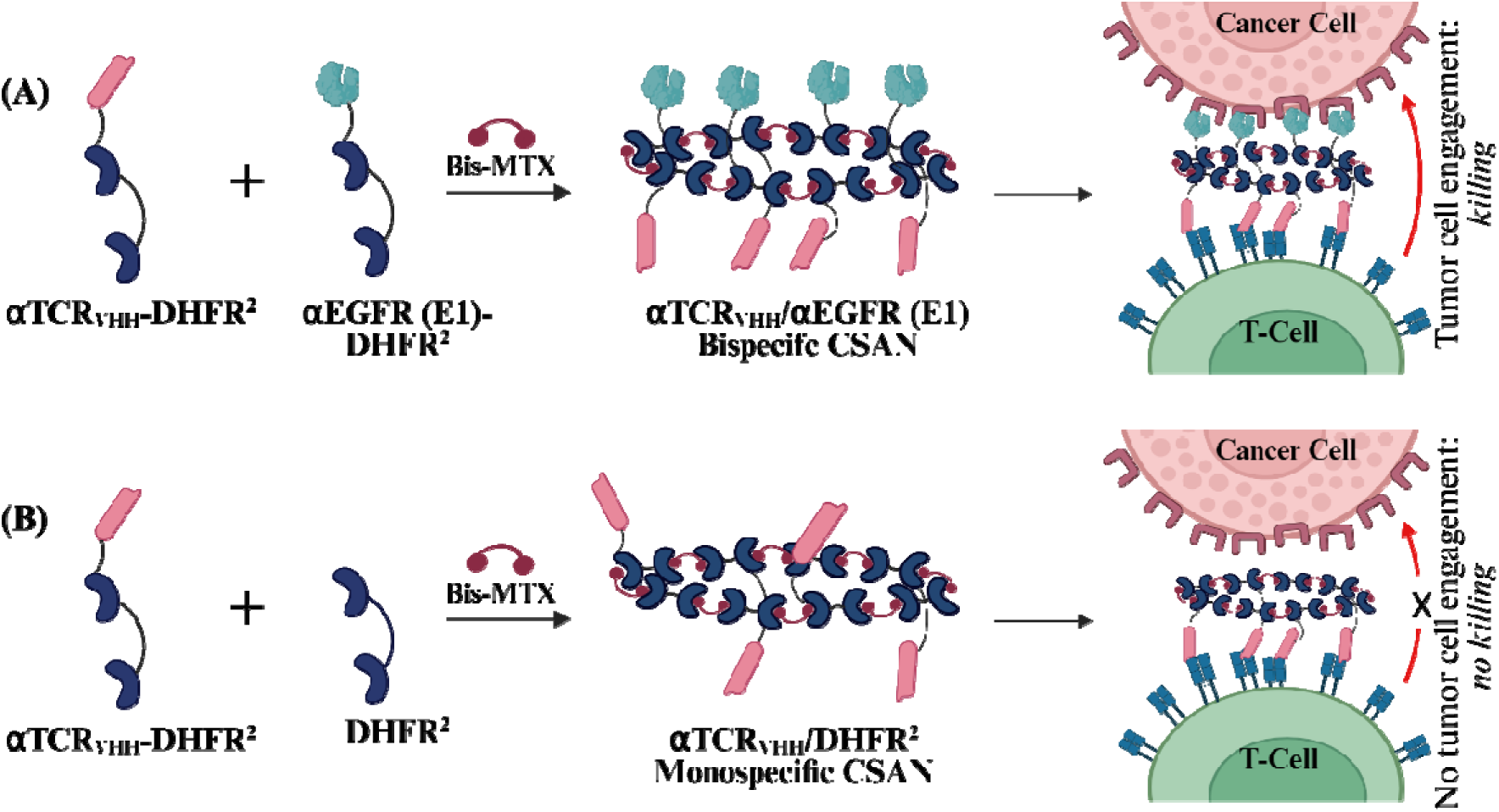
Schematic Representation of Chemically Self-Assembled Nanorings (CSANs) as T-Cell Engagers. Dimers of Dihydrofolate Reductase (DHFR^2^) spontaneously self-assemble to form CSANs upon addition of the chemical dimerizer, BisMTX. T-cell targeting domain (αTCR_VHH_) and tumor-targeting domain (αEGFR) are fused to the DHFR^2^ units to generate αTCR_VHH_-DHFR^2^ and DHFR^2^**/**αEGFR (E1) fusion proteins. (A) Equimolar amounts of αTCR_VHH_-DHFR^2−^ and DHFR^2^**/**αEGFR (E1) are mixed with BisMTX to generate multivalent bispecific αTCR_VHH_ /αEGFR (E1) CSANs that can cross-link T-cells and EGFR+ tumor cell leading to tumor cell death. (B) Non-targeted monospecific αTCR_VHH_ / DHFR^2^ CSANs lacking tumor-targeting domain cannot facilitate tumor-cell engagement and subsequent cell lysis.

Previously, our laboratory has demonstrated the efficacy of αCD3 scFv based CSANs against targets such as EpCAM, EGFR, CD133, and B7H3 both *in vitro* and *in vivo*^25–28^. However, due to the inherent challenges associated with the scFv scaffold targeting TCR/CD3 complex and safety concerns with the high affinity CD3 engagement, we sought to target the TCR directly as signals provided by its engagement has been demonstrated to be more physiological as compared to CD3 engagement^29^. While the exact signaling cascades initiated after engagement of TCR versus CD3 of the TCR/CD3 remains complex, the consequent functional difference is well documented^29–32^. Targeting different epitopes in the TCR/CD3 complex, as well as the strength of engagement, determines the degrees of T-cell activation, proliferation and cytokine release. Monoclonal antibodies against the TCR α/β such as BMA031 have demonstrated potent cytotoxicity with reduced systemic cytokine toxicity and thus a superior safety profile compared to classical CD3 binders such as OKT3.^33^ A recent study targeting AML with an αTCR-nanobody demonstrated effective tumor control without the severe cytokine release characteristic of CD3-targeted TCE.^34^ Therefore, after screening several candidates, we chose to utilize a nanobody (αTCR_VHH_) targeting the TCR α/β constant region in our CSAN platform. It is important to note that while engaging T-cells via the TCR subunit may initiate distinct intracellular signaling pathways compared to the highly explored CD3 engagement, a detailed dissection of these downstream pathways is beyond the scope of the current study. The primary objective of this study was to validate whether CSANs composed of αTCR_VHH_ and TAA targeting domain would maintain anti-tumor efficacy while widening the therapeutic window. We also wanted to address the overwhelming reliance on the CD3 subunit as the universal domain for T-cell engagement. By validating the efficacy of targeting a different subunit of the TCR/CD3 complex, we aim to diversify T-cell engagement epitopes. Establishing the TCR constant region as a viable and equally efficient alternative would expand the landscape for designing future immunotherapeutics, providing a diverse family of binding domains that can be explored for specific kinetic or safety profiles rather than being reliant on a singular target.

Through this work, we were able to demonstrate that CSANs composed of αTCR_VHH_ as T-cell binding domain successfully induced potent, dose-dependent cytotoxicity against solid tumor models expressing two different TAAs, EGFR and PSMA, thus showing applicability against a broad range of tumor targets. Furthermore, we showed that the activity is modulated by antigen density on the tumor cell-surface. Upon comparison with the CSANs that utilize αCD3_scFv_ as T-cell binding element, we showed that the binding affinity to the TCR/CD3 binding domain strongly influences the cytotoxic activity, activation of immune cells, specificity of action and safety against non-targeted toxicity. Taken together, these results demonstrate that nanobody based CSANs provide a robust, structurally simplified, safer alternative to scFv-based engagers, offering a versatile platform for the development of solid immunotherapies.

## 2. RESULTS AND DISCUSSION

### 2.1 Preparation and characterization of αTCR_VHH_-DHFR^2^ fusion proteins and CSANs

In this study, we systematically engineered a T-cell engager with a nanobody targeting the TCR α/β constant region (αTCR_VHH_) to overcome the inherent challenges associated with scFvs. In previous studies Roobrouck et. al. reported that an αTCR_VHH_ when fused with immunoglobulin single variable domains (ISVs) that target tumor associated antigens, induces T-cell activation and subsequent target cell lysis. Moreover, target cell lysis was observed only when the fusion proteins bridge the immune cell and target cell. Among the many clones reported, Clone T0170056G05 was chosen due to its high potential to induce cytotoxicity and T-cell activation against multiple tumor cell lines as a part of a bispecific fusion protein having a tumor targeting ISV.^35^

Structural analysis places the N-terminus of the V_HH_ near the 3^rd^ Complementarity Determining Region (CDR3), which plays a dominant role in epitope recognition and governs antigenic diversity and specificity. Any modification at the N-terminus could therefore compromise the integrity of the antigen-binding paratope through steric interference.^19, 36^ Guided by these structural constraints, we designed the αTCRV_HH_ construct with an unmodified N-terminus and fused the DHFR^2^ domain to the C-terminus of the nanobody to preserve binding functionality (Figure 2A).

**Figure 2.**
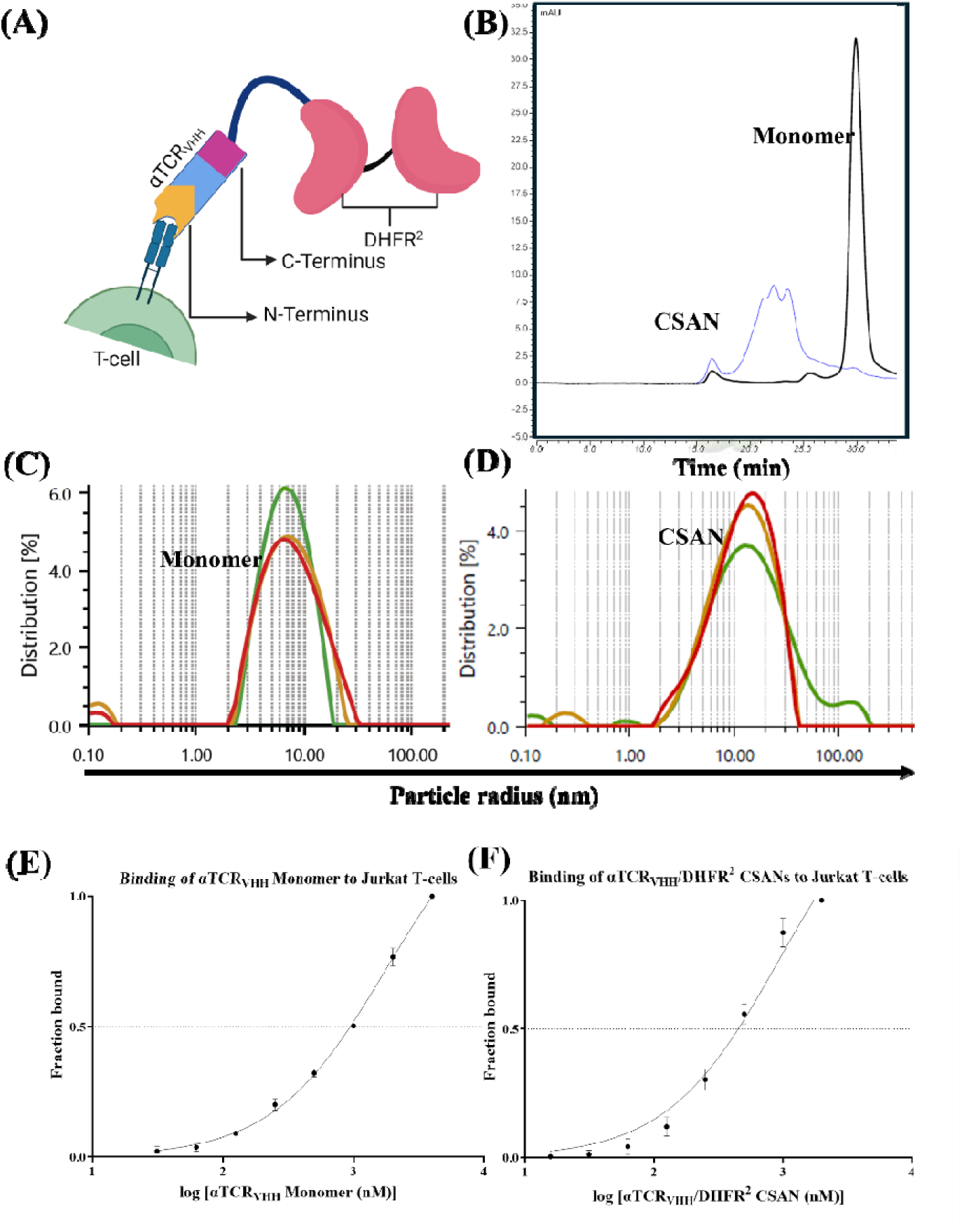
αTCR-DHFR^2^ Design and Characterization (A) Schematic representation of the protein monomer binding to the T-cell surface. (B) Size Exclusion Chromatography (SEC) confirmed the monomeric protein purity as well as the CSAN formation upon addition of Bis-MTX. The hydrodynamic radius of the monomer (C) and CSAN (D) was determined by Dynamic Light Scattering (DLS). Data is representative of 3 experiments. Flow cytometry was used to determine the binding affinity of the monomer (1.77 μM) and that of CSAN (0.93 μM) (E and F) using Jurkat T-cells.

Nanobodies have at least one disulfide bond linking the CDR3 to framework region (FR) or other CDR regions that are responsible for stabilization.^37^ Note here that due to the reductive environment of the cytoplasm, in traditionally used *E. coli* strains (BL21, Rosetta, etc.) the expression of nanobodies may undergo misfolding or loss of functionality because of the reduction of these disulfide bonds; thus, leading to expression as inclusion bodies and the need for highly cumbersome purification steps involving refolding of the protein. Periplasmic expression of proteins is well-known and favored for proteins with disulfide bonds due to the oxidative environment in the periplasm. However, various hurdles such as low yield, accumulation of protein in the cytoplasm, degradation of protein expressed in periplasm, etc. have been reported.^38^ ‘*T7 Express SHuffle Cells*’ are genetically engineered *E. coli* that promote disulfide bond formation in the cytoplasm. They are engineered to constitutively express disulfide bond isomerase DsbC that promote correction of mis-oxidized proteins into their correctly folded forms. This strain reduces the cytoplasmic reductive pathways, thereby allowing the formation of proper disulfide bonds.^39, 40^ Therefore, we chose ‘*T7 Express Shuffle Cells*’ as the expression system for αTCR_VHH_-DHFR^2^ fusion proteins. The protein was expressed as a soluble protein, which allowed easier and faster protein purification using Immobilized Metal Affinity Chromatography (IMAC) (See Figure S1A and S1B in S.I.). Observation of a single sharp peak at 30 min corresponding to the purified αTCR_VHH_-DHFR^2^ monomer in Size Exclusion Chromatography (SEC) confirmed the successful expression and purification of the protein (Figure 2B). SEC was also used to characterize the oligomerization of the CSANs composed of αTCR_VHH_-DHFR^2^ and DHFR^2^ proteins in the presence of the chemical dimerizer, bis-MTX, mixed in a 1:1:2.5 ratio and incubated at room temperature for 1 hour. We observed a lower retention time (∼ 23min, Figure 2B) for CSANs owing to their larger hydrodynamic radius compared to smaller monomers, which is consistent with previous reports of CSANs formation. Additionally, the oligomerization of αTCR_VHH_-DHFR^2^ in the presence of bis-MTX was observed by Dynamic Light Scattering (DLS) experiments, revealing an increase in the hydrodynamic radius from 6.78±0.27 nm (Figure 2C) (corresponding to αTCR_VHH_-DHFR^2^) to 11.73±1.33 nm (Figure 2D) (corresponding to DHFR^2^/DHFR^2^ CSANs). It is important to note that the polydispersity index of CSANs is within ∼ 20%, which is acceptable as being a uniform sample with respect to sample size. These observations confirm the functionality of the DHFR^2^ domain in the engineered αTCR_VHH_-DHFR^2^ fusion protein.

Next, we estimated the binding affinity (K_d_) on Jurkat T-cells to validate the antigen targeting functionality of the αTCR_VHH_-DHFR^2^ protein monomer and CSAN using flow cytometry. Jurkat T-cells were incubated with different concentrations of αTCR_VHH_-DHFR^2^ for 2 hours followed by a subsequent incubation with PE labeled αFLAG antibody, which binds to the FLAG tag on the αTCR_VHH_-DHFR^2^ protein. The binding affinity of the monomer and CSANs was determined by measuring the PE fluorescence of the antibody associated with the Jurkat T-cell bound αTCR_VHH_-DHFR^2^. The K_d_ of the monomeric αTCR_VHH_-DHFR^2^ protein for the Jurkat cells was found to be 1.77 μM (Figure 2E). Upon assembly into the oligomeric CSANs, the K_d_ decreased to 0.93 μM (Figure 2F). This observed reduction of K_d_ reflects the avidity effect central to the CSAN architecture, with multivalency improving the overall binding affinity to the cell surface. Previously designed DHFR^2^-αCD3_scFv_, which binds to the CD3ε chain of the TCR/CD3 complex on the T-cells was used as a positive control for T-cell binding and activation.^23, 41^ Consistent with previous reports, the scFv constructs exhibited stronger binding affinity.^41^ For cancer cell targeting, the previously reported αEGFR (E1)-DHFR^2^ protein was employed, in which the EGFR-binding fibronectin domain E1 is fused to DHFR^2^ at its C-terminus.^27^ The expression and purification of DHFR^2^-αCD3 and αEGFR E1-DHFR^2^ were carried out according to previously reported protocols.^27^

### 2.2 áTCR_VHH_ /αEGFR (E1) CSANs are selectively cytotoxic to EGFR positive cell lines *in vitro*

Antigen density on tumor cell surfaces is a key determinant of efficient immunological synapse formation and the subsequent T-cell engager-mediated cytolysis of tumor cells.^42^

To determine the effect of antigen expression on cytotoxicity, we assessed the cytotoxicity of αTCR_VHH_ /αEGFR (E1) CSANs on 3 different cell lines with distinct EGFR expression profiles: A431-R (∼10^6^ receptors/cell), MDA-MB-231-G (∼68000 receptors/cell) and MCF-7-R (∼5000 receptors/cell). The cytotoxicity of the TCR targeted aEGFR CSANs was compared against αCD3_scFv_/αEGFR (E1) CSANs. Monospecific CSANs lacking the tumor-targeting domain αEGFR (E1) (αTCR_VHH_/DHFR^2^, and αCD3_scFv_/DHFR^2^) were included as controls to determine the T-cell activating ability of the CSANs without engaging the tumor cells. CSAN mediated T-cell cytotoxicity was investigated on a monolayer of tumor cells co-cultured with T-cells freshly isolated from human PBMCs and treated with variable concentrations of the CSANs over a 72-hour period. Cytotoxicity at variable Effector to Target Cell (E:T) ratios was investigated to assess the intrinsic T-cell engagement potency of the CSANs toward the target EGFR positive tumor cells.^43^ In a co-culture of A431-R cells, αTCR_VHH_ /αEGFR (E1) CSANs exhibited potency comparable to the αCD3_scFv_/αEGFR (E1) CSANs at high E:T ratios (Figure 3A and Figure S2A). At E:T ratio of 10:1, both constructs achieved maximal cytotoxicity at the highest CSAN concentration. However, as the CSAN concentration decreased, the αCD3_scFv_/αEGFR (E1) CSANs maintained moderate potency, whereas the cytotoxicity of the αTCR_VHH_ variant underwent a steeper declined (Figure 3B and Figure S2B). This observation suggests that the higher binding affinity of the DHFR^2^-αCD3_scFv_ facilitates T-cell activation and tumor cell engagement at lower concentrations than DHFR^2^-αTCR_VHH_. Cytotoxicity toward the tumor cells was found to be dependent on tumor cell targeting since neither the αCD3 nor the αTCR monospecific CSANs induced tumor cell cytotoxicity.

**Figure 3.**
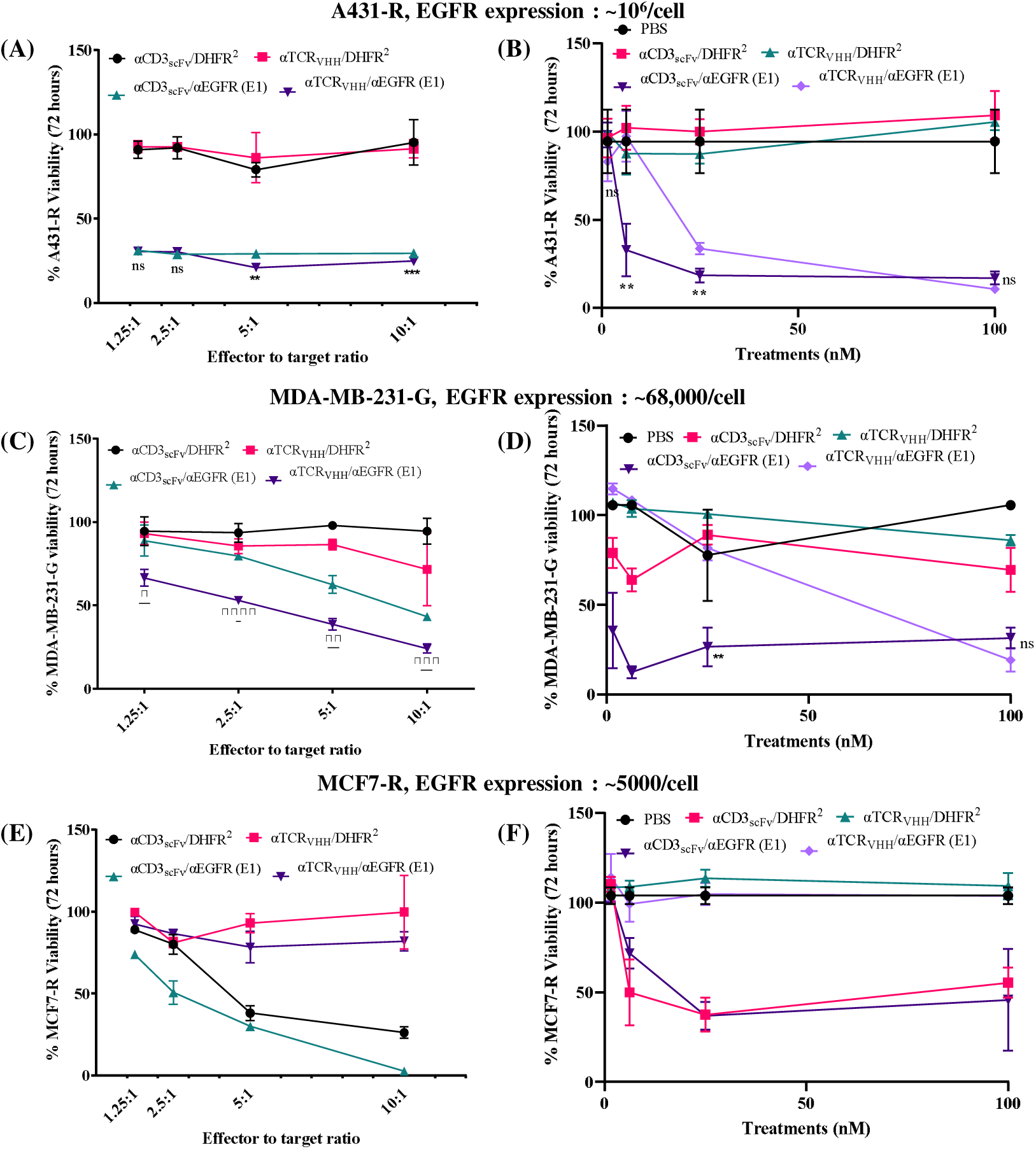
αTCR_VHH_/αEGFR (E1) Bispecific CSANs direct cytotoxicity against EGFR+ cell lines. Tumor cells were seeded in 96-well plate as a monolayer. T-cells freshly isolated from healthy donor PBMCs were added to the wells 20 hours after with monospecific or bispecific CSANs. Tumor cell viability was monitored over 72 hours. Each bullet point represents the final tumor cell count at the end of 72 hour cytotoxicity assay. A431-R cell viability (A), MDA-MB-231-G cell viability (C), and MCF7-R cell viability (E) were monitored over 72 hours at a fixed CSAN concentration (100nM) and different E:T ratio. End point cell viability data at each E:T ratio normalized with respect to treatment with PBS only is shown. Significance of αTCR_VHH_/αEGFR (E1) was calculated with respect to of αCD3scFv/αEGFR (E1) by two tailed unpaired t-test. Effects of different concentrations of CSANs on the viability of A431-R cells (B), MDA-MB-231-G cells (D) and MCF7-R cells (F) were analyzed at a 5:1 E:T ratio. All data is normalized to tumor cells not treated with T-cells or CSANs. Significance of αTCR_VHH_/αEGFR (E1) was calculated with respect to αCD3_scFv_/αEGFR (E1) by 2-tailed Student’s *t* test. (Data is displayed as mean ± SD, *P<0.03, **P<0.007, ***P<0.0006, ****P<0.0001). Data shown is obtained from one donor but is representative of two donors (See Figure S2 in S.I.)

The divergence in potency became more pronounced in MDA-MB-231-G cells (moderate EGFR expression). While comparable cytotoxicity was seen for both bispecific CSANs at higher E:T ratios and high concentrations, there was a more pronounced reduction of potency for αTCR_VHH_/αEGFR (E1) CSANs at lower T-cell density and lower concentrations than for the αCD3_scFv_/αEGFR (E1) CSANs (Figure 3C, 3D, Figure S2C and S2D). Notably, the monospecific αCD3_scFv_ CSANs elicited observable cytotoxicity even without tumor-cell engagement, indicating a higher propensity for non-specific activity. In contrast, the monospecific αTCR_VHH_ CSANs construct triggered no off-target cytotoxicity.

The difference in selectivity was most evident for MCF7-R cells (negligible/low EGFR expression). The αCD3_scFv_ CSANs induced significant cytotoxicity despite the lack of significant antigen expression, and killing was observed even with the non-targeting monospecific controls. Conversely, the αTCR_VHH_ CSANs showed no cytotoxicity towards MCF7-R cells, even at the highest E:T ratio (10:1) (Figure 3E). However, this observation is subject to the inherent heterogeneity in the T-cells from different donors, where both the constructs showed comparable cytotoxicity at 10:1 E:T. The αTCR_VHH_ CSANs still maintained a better safety margin, with significant decline in cytotoxicity at lower E:T ratios (See Figure S2E in S.I.). The αTCR_VHH_ CSANs also induced significantly less cytotoxicity compared to αCD3_scFv_ CSANs at the CSAN concentrations tested across different donors (Figure 3F and Figure S2F).

Taken together, these results indicate that the weaker-binding αTCR_VHH_ CSANs require robust engagement with the TAAs to generate a strong immune synapse and elicit a cytotoxic response. While the αCD3_scFv_ constructs displayed higher potency at lower concentrations, significant cytotoxicity was observed by the αTCR_VHH_ CSANs regardless of the level of EGFR expression by the tumor cells.

### 2.3 Effect of αTCR_VHH_/αEGFR (E1) CSANs on T-cell activation against EGFR^HIGH^ (A431-R) and EGFR^MODERATE^ (MDA-MB-231-G) cell line

To assess whether the lower binding affinity of the αTCR_VHH_ alters the quality of T-cell activation, we conducted a comprehensive profiling of T-cell activation markers from the co-cultures with A431-R (EGFR^HIGH^) and MDA-MB-231-G (EGFR^MODERATE^) cell lines. We determined the expression level of early (CD69), and late (CD25) activation marker along with the T-cell exhaustion marker (PD-1) on both CD4+ and CD8+ T-cell subsets across a time range of 24 hours and 72 hours.

In the EGFR^HIGH^ (A431-R) model, the activation markers revealed a clear distinction in the kinetics of T-cell activation by the αTCR_VHH_ binder versus the αCD3_scFv_ counterpart. At the early 24-hour time point, T-cells stimulated by the bispecific αTCR_VHH_/αEGFR (E1) CSANs exhibited significantly lower expression of CD69 compared to the higher affinity αCD3_scFv_/αEGFR (E1) CSANs (Figure 4A, 4G, Figure S3A and S3G). However, by 72 hours, this trend was reversed: i.e., CD69 expression by αTCR_VHH_/αEGFR (E1) CSANs was significantly higher than the αCD3_scFv_ counterpart (Figure 4D, 4J, Figure S3D and S3J). We postulate that this trend reflects the slower activation kinetics for the αTCR_VHH_/αEGFR (E1) CSANs relative to the αCD3_scFv_/αEGFR (E1) CSANs. Owing to its lower K_d_, the receptor occupancy by the αCD3_scFv_/αEGFR (E1) CSANs is likely higher than the αTCR_VHH_/αEGFR (E1) CSANs at equivalent concentrations. Consequently, the αTCR_VHH_/αEGFR (E1) CSANs require a longer time period to form a stable immune synapse and reach the appropriate signaling threshold. This dynamic could also explain the delay in CD69 expression suggesting a slower ramping up of T-cell activation when the TCR is targeted compared to CD3. The monospecific αCD3_scFv_ CSANs (lacking E1: the tumor targeting domain) induced robust upregulation of CD69 on both CD4+ and CD8+ T-cells, even in the complete absence of tumor cells (Figure 4 and Figure S3), reflecting ‘*tonic signaling*’. In contrast, the monospecific αTCR_VHH_ CSANs induce negligible CD69 expression, thus the concentration required for αTCR_VHH_ CSANs induced “tonic signaling” is significantly higher, requiring tumor engagement to initiate signaling at lower concentrations than αCD3_scFv_/αEGFR (E1) CSANs.

**Figure 4.**
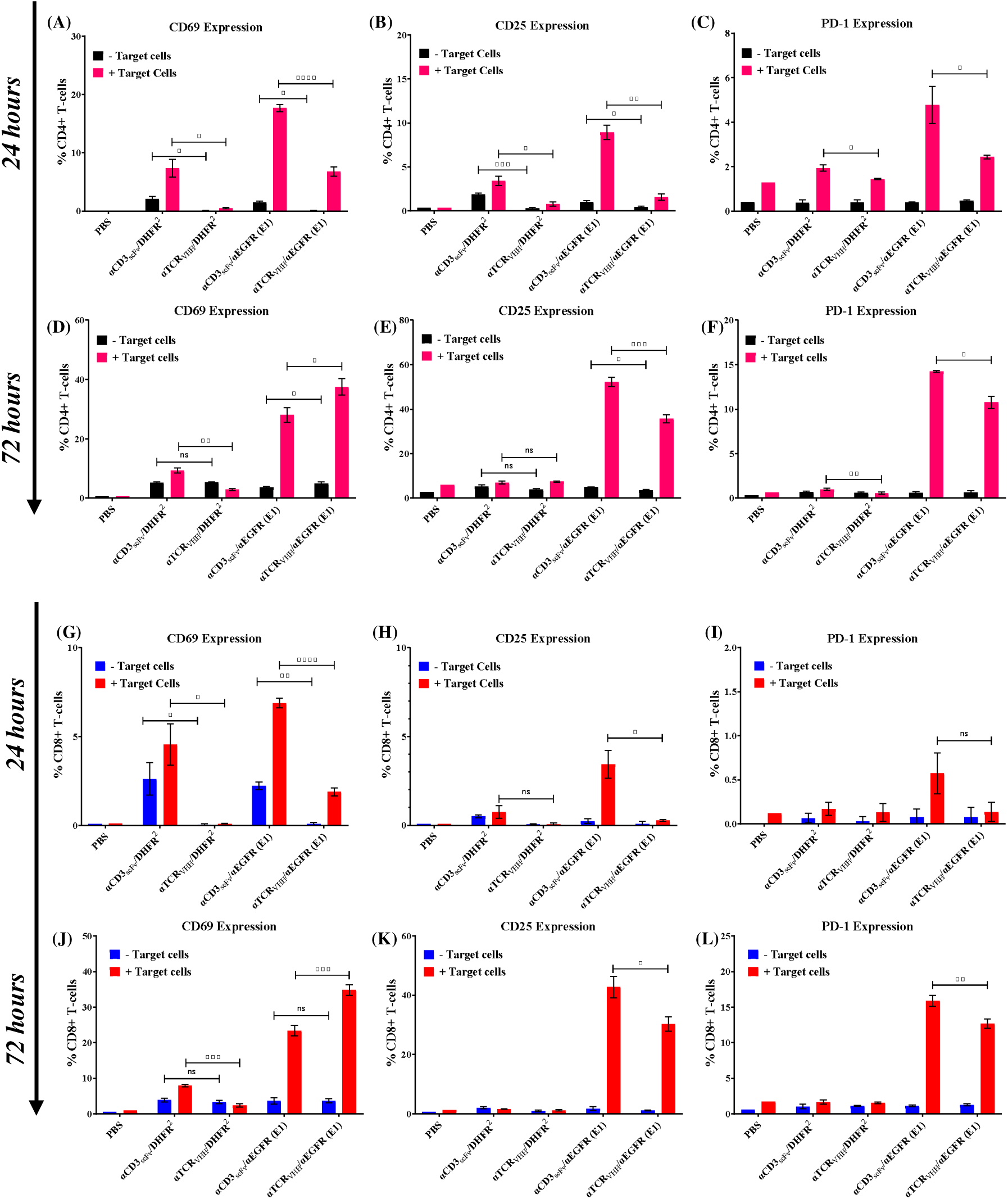
αTCR_VHH_/αEGFR (E1) Bispecific CSANs activate T-cells selectively in cytotoxicity assay with A431-R cells at 10:1 E:T ratio with 100nM of each treatment. T-cells isolated from healthy donor PBMCs were co-cultured with media, monospecific or bispecific CSANs in presence or absence of A431-R cells. CD69 expression on CD4+ and CD8+ T cells were measured at 24 hours (A,G) and at 72 hours (D,J). CD25 expression on CD4+ and CD8+ T-cells were measured at 24 hours (B,H) and at 72 hours (E,K). PD-1 expression on CD4+ and CD8+ T-cells were measured at 24 hours (C,I) and at 72 hours (F,L). Data shown is obtained from one donor but is representative of two donors (See Figure S3 in S.I.). Significance for different treatments is calculated by 2-tailed Student’s *t* test. (Data is displayed as mean ± SD, *P<0.07, **P<0.0095, ***P<0.0008, ****P<0.0001).

Analysis of the high affinity IL-2 receptor, (CD25: late activation marker), revealed a notable functional uncoupling between the surface phenotype of activated T-cells and their cytotoxic capacity. In the A431-R model, the αTCR_VHH_/aEGFR (E1) CSANs induced significantly lower CD25 expression than αCD3_scFv_/aEGFR (E1) CSANs at 24, and 72 hours (Figure 4B, 4H, Figure S3B and S3H) and (Figure 4E, 4K, Figure S3E and S3K). Despite the slower rate of full activation, the end-point cytotoxicity was comparable between the two groups. These results indicate that maximal induced target cytotoxicity can be reached even with the slower rate of T-cell activation by αTCR_VHH_/αEGFR (E1) CSANs to T-cells, compared to αCD3_scFv_/αEGFR (E1) CSANs.

Prolonged hyper-stimulation of T-cells is known to promote dysfunctional and exhausted state. To evaluate this effect, we quantified the expression of PD-1, an immune checkpoint receptor whose upregulation, upon engagement with PD-L1 on tumor cells, is closely associated with T-cell exhaustion and immune evasion.^44^ At both 24 and 72 hours, T-cells stimulated by αTCR_VHH_/αEGFR (E1) CSANs exhibited significantly lower expression of PD-1 compared to αCD3_scFv_/αEGFR (E1) CSANs (Figure 4C, 4F, 4I, 4L, Figure S3C, S3F, S3I and S3L). This reduced expression of the checkpoint inhibitor on the T-cells suggests that the attenuated signaling by the nanobody-based CSAN may preserve the effector capabilities of the T-cells over a longer duration without driving them into an exhausted state, as is common for higher-affinity T-cell engagers.^45^

When either αTCR_VHH_/αEGFR (E1) CSANs or αCD3_scFv_/αEGFR (E1) CSANs were used to target T-cells to the EGFR^MODERATE^ (MDA-MB-231-G) cell line, T-cell activation was found to be faster when targeting CD3 than the TCR, with a few exceptions. Unlike targeting EGFR^HIGH^ A431-R cells where CD25 peaked late (72hours), T-cells co-cultured with EGFR^MODERATE^ MDA-MB-231-G cells showed accelerated activation, with >50% of T-cells expressing CD25 by 24 hours for both the nanobody and scFv based bispecific constructs (Figure 5B, 5H, Figure S4B and S4H). While CD25 expression for T-cells stimulated by αTCR_VHH_/αEGFR (E1) CSANs initially lagged behind the αCD3_scFv_/αEGFR (E1) CSANs, this distinction was lost by 72 hours (Figure 5E, 5K, Figure S4E and S4K).

**Figure 5.**
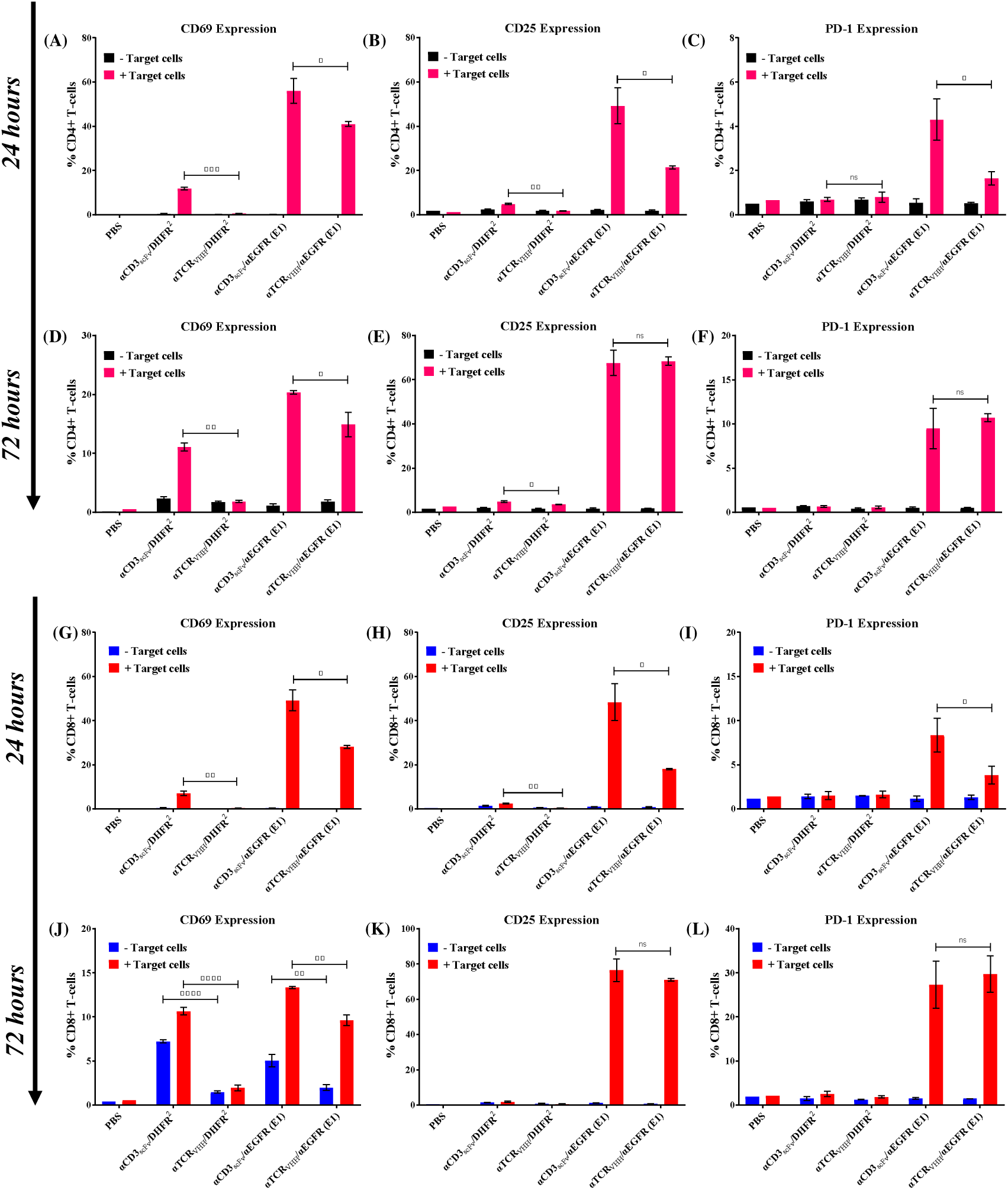
αTCR_VHH_/αEGFR (E1) Bispecific CSANs activate T-cells in cytotoxicity assay with MDA-MB-231-G cells 5:1 E:T ratio with 100nM of each treatment. T-cells isolated from healthy donor PBMCs were co-cultured with media, monospecific or bispecific CSANs in presence or absence of MDA-MB-231-G cells. CD69 expression on CD4+ and CD8+ T cells were measured at 24 hours (A,G) and at 72 hours (D,J). CD25 expression on CD4+ and CD8+ T-cells were measured at 24 hours (B,H) and at 72 hours (E,K). PD-1 expression on CD4+ and CD8+ T-cells were measured at 24 hours (C,I) and at 72 hours (F,L). Data shown is obtained from one donor but is representative of two donors (See Figure S4 in S.I.). Significance for different treatments is calculated by 2-tailed Student’s *t* test. (Data is displayed as mean ± SD, *P<0.05, **P<0.009, ****P<0.0001).

The higher expression CD25 after treatment with either αTCR_VHH_/αEGFR (E1) CSANs or αCD3_scFv_/αEGFR (E1) CSANs and higher initial expression of CD69 is consistent with a higher level of T-cell activation for both CD4+ and CD8+ T-cells in the presence of EGFR^MODERATE^ MDA-MB-231-G than EGFR^HIGH^ A431-R cells. The higher level of T-cell activation did not result in a significant increase in CD4+ T-cell PD-1 expression; in contrast to CD8+ T-cells, in which a higher level of PD-1 expression was observed for treatment with both αTCR_VHH_/αEGFR (E1) CSANs or αCD3_scFv_/αEGFR (E1) CSANs. One potential explanation for the discrepancy between the effect of the target cells on the T-cells by EGFR^MODERATE^ MDA-MB-231-G and EGFR^HIGH^ A431-R cells may reside in the higher amount of ICAM expression by MDA-MB-231 cells over A431 cells.^46–48^ The higher expression by MDA-MB-231 may increase the probability of productive immune synapse formation, despite the lower level of EGFR expression.

The formation of memory T-cells is an important component of a strong, sustained anti-tumor immune response and hence a critical goal for TCE development. To evaluate this, we determined the differentiation of CD4+ and CD8+ T-cells into central memory (CD45RO^+^/CCR7^+^) and effector memory (CD45RO^+^/CCR7^−^) subsets after 72 hours of co-culture with A431-R and MDA-MB-231-G cells in presence of the different CSANs. While both αCD3_scFv_/αEGFR (E1 and αTCR_VHH_/αEGFR (E1) CSANs generated memory T-cell populations in some donors, the heterogeneity of T-cells isolated from different donor PBMCs led to an indefinite conclusion. Because the responses varied across different donors, we were unable to establish a statistically robust pattern regarding how the different binding epitopes and affinities of the two types of CSANs affect the generation of memory T-cells (See Figure S5 and S6 in S.I.).

### 2.4 Effect of αTCR_VHH_/αEGFR CSANs on Regulatory T-cell formation against EGFR^HIGH^ (A431-R) and EGFR^MODERATE^ (MDA-MB-231-G) cell line

A critical challenge in the recruitment of T-cell engagers is the inadvertent activation and recruitment of regulatory T-cells (T_regs_) While T_regs_ are essential for the maintenance of immune homeostasis and prevention of over-inflammation of the immune system, their accumulation in the tumor microenvironment (TME) contribute to the immunosuppressive environment and pose a significant barrier to immunotherapy.^49, 50^ T_regs_ polarize the immune system and dampen anti-tumor responses through the secretion of inhibitory cytokines (e.g. IL-10, TGF-β) suppressing functions of CD4, CD8, NK, and NK-T cells, cytolysis of effector T cells and NK cells and impairment of APCs such as DCs.^51^ Since conventional T-cell engagers can provide the strong stimulus required to activate and expand this immunosuppressive population, we sought to determine if the reduced binding affinity of the αTCR_VHH_-based CSANs could mitigate this effect. We employed flow-cytometry to monitor the activation and expansion of T_regs_ following treatment of EGFR^HIGH^ (A431-R) and EGFR^MODERATE^ (MDA-MB-231-G) cell lines. T_regs_ are defined as a subset of CD4+ T-cells exhibiting high CD25 and low to negative CD127 (CD4+/CD25+/CD127^low/negative^), a surface phenotype that is inversely correlated with the master regulator FoxP3.^52, 53^

The high affinity construct (DHFR^2−^αCD3_scFv_/ αEGFR (E1) induced T_reg_ expansion in both the cell lines after 72 hours (Figure 6A and 6B), which is found to be consistent with the hypothesis that T-cell signaling strength dictates T_regs_ proliferation. Conversely, the treatment with αTCR_VHH_/αEGFR (E1) CSANs resulted in significantly lower T_reg_ expansion. Interestingly, the extent of T_reg_ expansion is also dependent on the intrinsic T-cell heterogeneity of different donors. The impact of signaling strength on the extent of T_reg_ expansion seems to rely on the immune profile and activation kinetics of individual donors (See Figure S7 in S.I.). Notably, in the MDA-MB-231-G cell line, even the monospecific DHFR^2^-αCD3_scFv_/DHFR^2^ CSANs lead to a significantly higher activation and expansion of regulatory T-cells compared to the αTCR_VHH_ construct (Figure 6B). Consistent with the effect seen on T-cell proliferation and activation (*vide supra*), when the amount of T_regs_ proliferation induced by the targeted T-cells interacting with the EGFR^HIGH^ expressing A431-R cells is compared to the EGFR^MODERATE^ expressing MDA-MB-231-R cells, approximately two-fold more T_regs_ were observed, suggesting that another factor other than EGFR expression is responsible for this difference.

**Figure 6.**
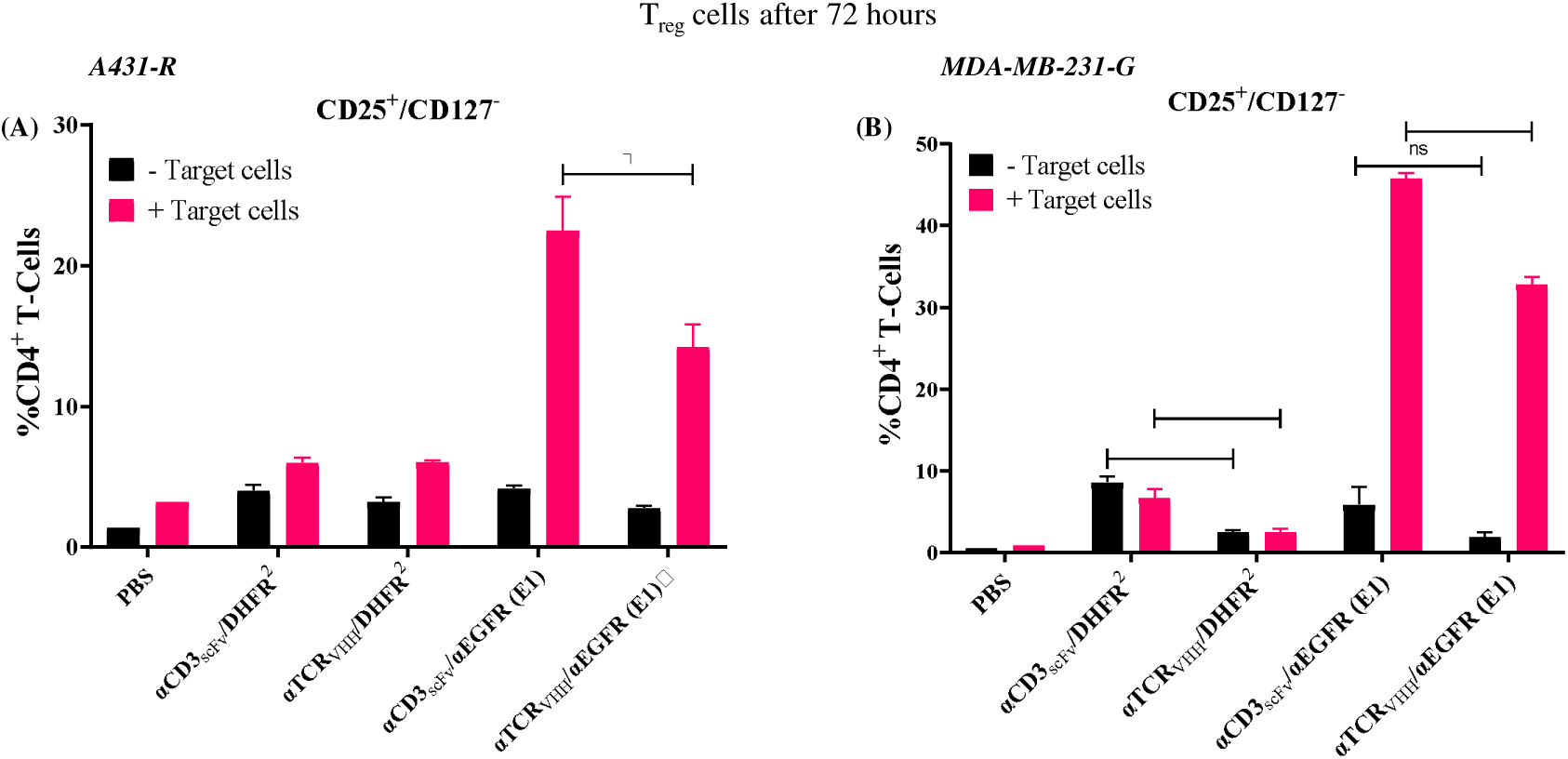
Regulatory T-cell response to CSANs *in vitro*. T-cells freshly isolated from healthy donor PBMCs were co-cultured with media, monospecific or bispecific CSANs in presence or absence of a 2D monolayer of A431-R (A) and MDA-MB-231-G (B) cells. Following 72 hour incubation, T-cells were harvested and analyzed for CD4, CD25 and CD127 expression. Significance was calculated by using 2-tailed Student’s *t* test. (Data is displayed as mean ± SD, *P<0.02, **P<0.003, ***P<0.0001) Data shown is obtained from one donor but is representative of two donors (See Figure S7 in S.I.).

These results suggest that the binding affinity of the T-cell engaging domain of the CSANs may potentially be an important determinant of immunosuppressive feedback. Thus, the αTCR_VHH_ CSANs may trigger cytotoxicity but with a reduced level of T_reg_ activation and proliferation compared to the αCD3_scFv_ depending on the inherent distribution of T-cell population in donors.

### 2.5 Effect of αTCR_VHH_ /αEGFR (E1) CSANs on cytokine release *in vitro*

*In vitro* release of cytokines by T-cells is a measure of the degree of activation.^54^ To correlate T-cell mediated cytotoxicity with immune activation and assess the potential for cytokine-related toxicity, we determined the release of Th1 cytokines in the co-culture supernatants at 24 and 72 hours for αTCR_VHH_/αEGFR (E1) CSANs and αCD3_scFv_/αEGFR (E1) CSANs.

Consistent with the established sequential release of cytokines upon T-cell activation, a temporal shift in the cytokine secretion for the αTCR_VHH_/αEGFR (E1) CSANs and αCD3_scFv_/αEGFR (E1) CSANs was observed. TNF-α is an early mediator of apoptosis, chemokine regulation and lymphocyte infiltration and reaches its peak at 12-24 hours following T-cell stimulation, which is then followed by the robust and sustained production of IFN-γ.^55–57^ This is reflected by the high levels of TNF-α release stimulated by both the bispecific CSANs at 24 hours (Figure 7A and Figure S9A), which then declined by 72 hours (Figure 7C and Figure S9C). Conversely, IFN-γ levels progressively increased from 24 hours to 72 hours, reflecting sustained effector functions (Figure 7B, 7D, Figure S9B and S9D).

**Figure 7.**
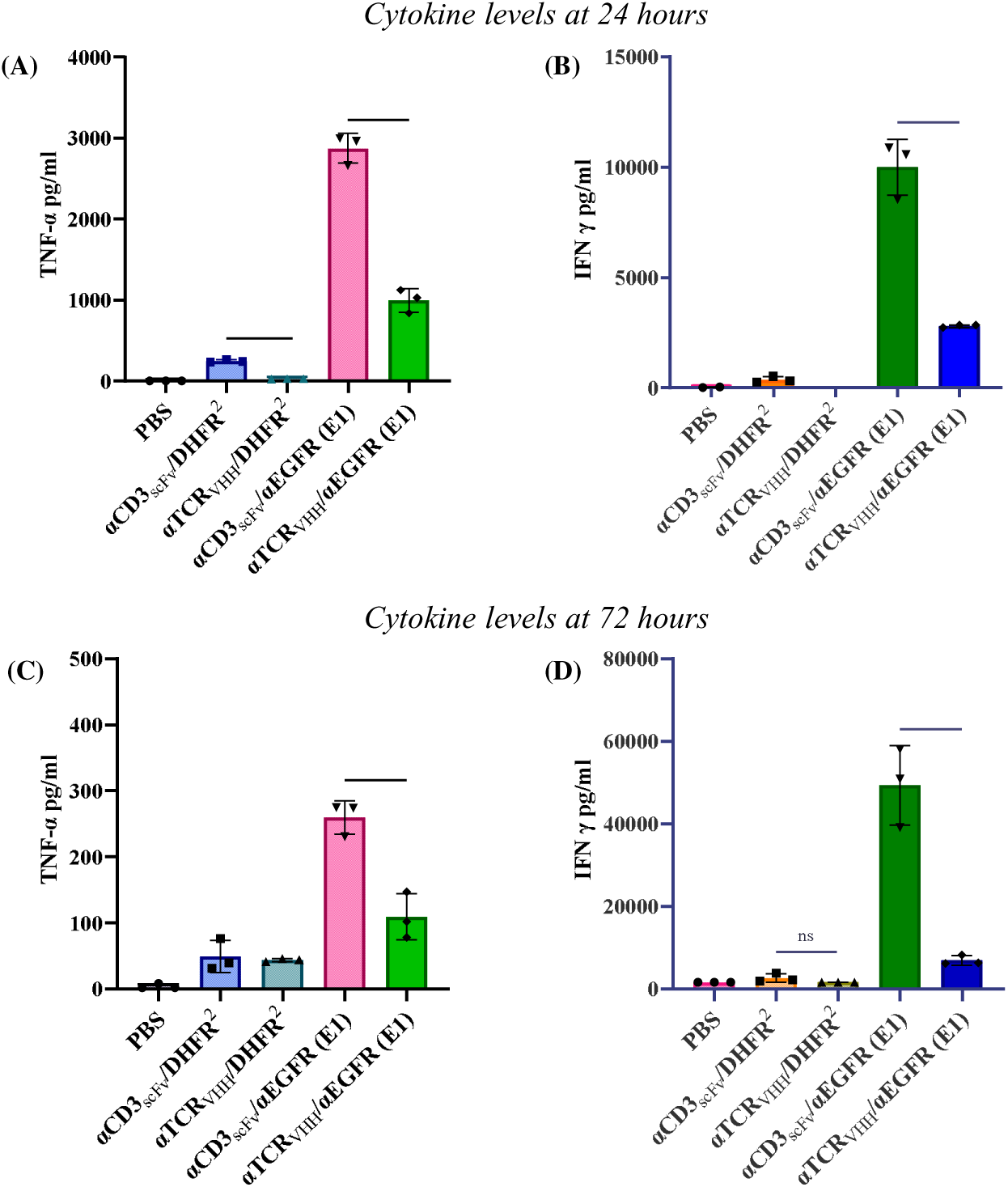
Determination of *in vitro* cytokine release as a result of αTCR_VHH_/αEGFR (E1) mediated MDA-MB-231-G cell lysis by T-cells freshly isolated from healthy donor PBMCs at 5:1 E:T ratio and 100nM treatments. Supernatants from the co-culture were analyzed for TNF-α (A) and IFN-γ (B) at 24 hours and 72 hours (C and D) using a sandwich ELISA. Significance was calculated using 2-tailed Student’s *t* test. (Data is displayed as mean ± SD, *P<0.02, **P<0.006, ***P<0.0003). Data shown is obtained from one donor but is representative of two donors (See Figure S9 in S.I.).

Stimulation strength of the TCR/CD3 complex dictates the magnitude of cytokine release which is consistent with the significantly higher levels of both TNF-α and IFN-γ by the stronger binding αCD3_scFv_/αEGFR (E1) CSANs compared to the αTCR_VHH_/αEGFR (E1) CSANs.^58^ Compared to the monospecific αCD3_scFv_ CSANs and αTCR_VHH_ CSANs controls, cytokine release was dependent on engagement with the tumor cells, with higher levels of TNFα being released at 24 h by the αCD3_scFv_ CSANs (Figure 7A, 7B and Figure S9).

### 2.6 áTCR_VHH_/αEGFR (E1) CSANs facilitate cytotoxicity in 3D cell culture

While 2D cell culture is a simple and cost-effective system to screen for cytotoxicity *in vitro*, it is overly simplified as it lacks the cellular organization and physical barriers inherent to solid tumors. 3D spheroids offer a higher degree of complexity, more closely mirroring the cellular organization and cell-cell interactions in solid tumors and hence are powerful tools to determine the efficacy of treatments.^59^ To validate the therapeutic potential of the CSAN constructs in a more physiologically relevant environment, we evaluated the cytotoxicity of αTCR_VHH_/αEGFR (E1) CSANs on 3D spheroids of EGFR^HIGH^ A431-R cells.

Spheroids were co-cultured with freshly isolated T-cells from healthy donor PBMCs (10:1 E:T ratio) and treated with bispecific CSANs or corresponding controls and the integrity of the spheroid monitored for 72 hours (Figure 8 and Figure S10). Spheroids treated with PBS or mono-specific controls retained their integrity and size. Treatment with the high affinity αCD3_scFv_/αEGFR (E1) CSANs resulted in a gradual reduction of the spheroid area, indicating effective surface-level killing, however, the core of the spheroids remained largely intact at the end point.

**Figure 8.**
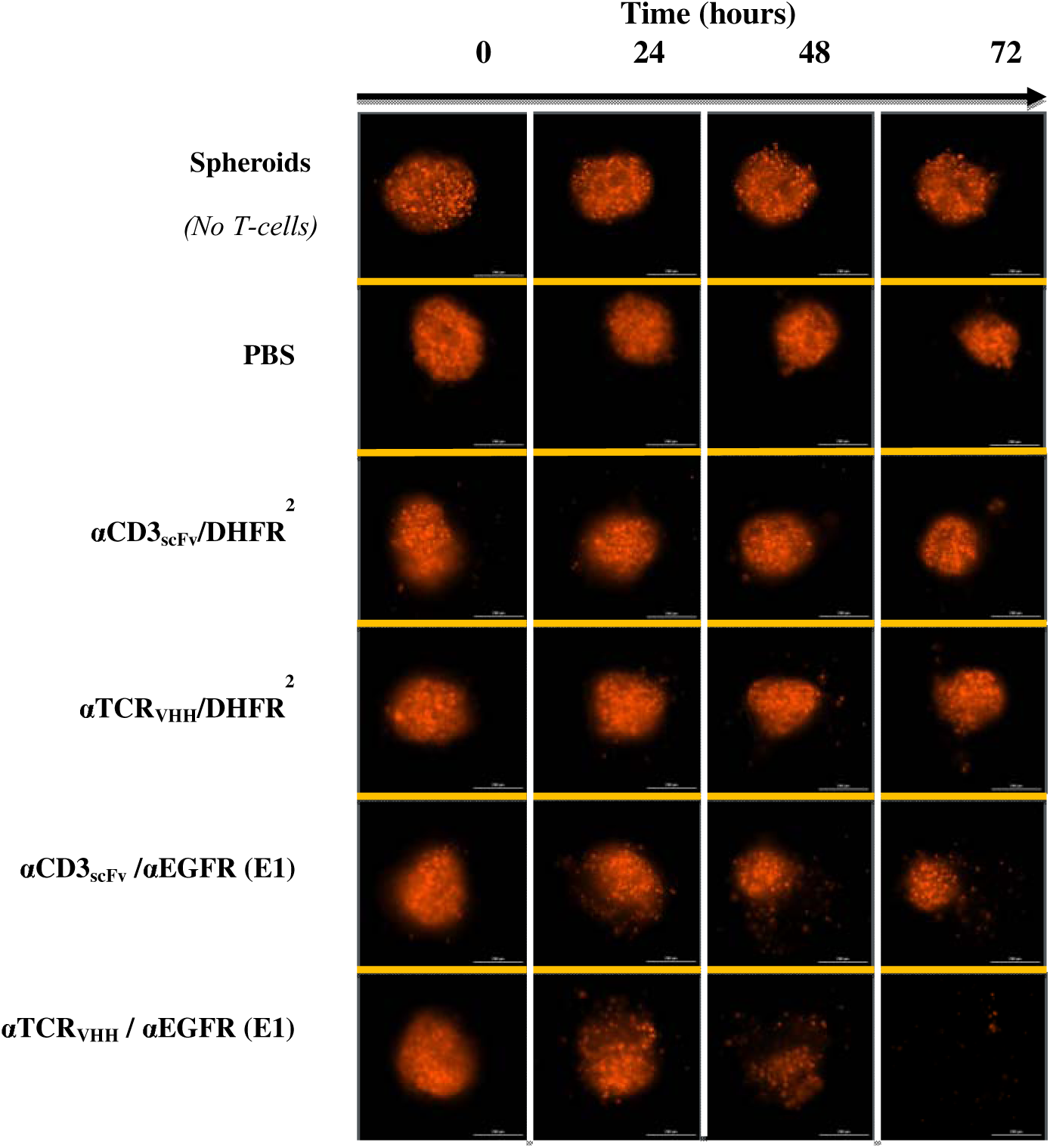
αTCR_VHH_/αEGFR (E1) Bispecific CSANs induced cytotoxicity against 3D spheroids of A431-R cells. Spheroids were incubated with freshly isolated T-cells from healthy donor PBMCs at 10:1 E:T ratio along with PBS, monospecific or bispecific CSANs at 100nM for 72 hours at 37°C and images were taken every 24 hours. Representative confocal images of the spheroids at different times points. Data shown is obtained from one donor but representative of two donors (See Figure S10 in S.I.).

In contrast, spheroids treated with αTCR_VHH_/αEGFR (E1) CSANs exhibited a more rapid and profound morphological disruption. By 72 hours, these spheroids had lost their structural integrity and undergone complete disintegration, outperforming the αCD3_scFv_ counterpart. The enhancement of spheroid cytotoxicity by the αTCR_VHH_/αEGFR (E1) CSANs compared to the αCD3_scFv_/αEGFR (E1) CSANs may reflect the lower affinity of the αTCR_VHH_ facilitating migration of the T-cells into the spheroid, which would not be observed in 2D cell culture.^60, 61^ Interestingly, this affinity dependent T-cell migration was not absolute and appeared to be donor dependent. For one specific donor, we observed no significant distinction in the 3D cytotoxicity between the high affinity and moderate-affinity CSANs. This indicates that the affinity-driven T-cell infiltration can be overridden by inherent T-cell motility and activation kinetics. Still, the monospecific αCD3_scFv_ CSANs showed slightly higher 3D cytotoxicity over the monospecific αTCR_VHH_ CSANs, consistent with our observations from the 2D assays (See Figure S10 in S.I.).

### 2.7 Evaluation of αTCR_VHH_ based CSANs targeting PSMA-expressing prostate tumor models

Having established the mechanistic advantages of αTCR_VHH_-CSANs in EGFR positive tumor models, we sought to explore the generalizability of the findings with another antigen. We chose Prostate Specific Membrane Antigen (PSMA) since it is a well-established biomarker for prostate cancer.^62, 63^ We evaluated the T-cell targeted cytotoxicity of αTCR_VHH_/αPSMA CSANs and αCD3_scFv_/αPSMA CSANs against the prostate cancer cell lines, C4-2-R (PSMA^POSITIVE^) and DU145-R (PSMA^NEGATIVE^). As observed for the EGFR targeted bispecific CSANs, the αTCR_VHH_/αPSMA CSANs were slower to induce targeted T-cell cytotoxicity to the C4-2-R (PSMA^POSITIVE^) cells than the αCD3_scFv_/αPSMA CSANs. Nevertheless, a similar level of cytotoxicity was achieved by 72 hours (Figure 9A, 9B, Figure S11A and S11B). However, unlike the breast cancer cells (with the exception of MCF-7 cells), non-targeted cytotoxicity by monospecific αCD3_scFv_ CSANs was significantly greater than observed for monospecific αTCR_VHH_ CSANs. When tested against the DU145-R PSMA^NEGATIVE^ cells, neither the monospecific nor the bispecific CSAN composed of either the αTCR_VHH_ or αCD3_scFv_ targeting domain induced cytotoxicity, even at the highest concentration and E:T ratios (Figure 9C, 9D, Figure S11C and S11D).

**Figure 9.**
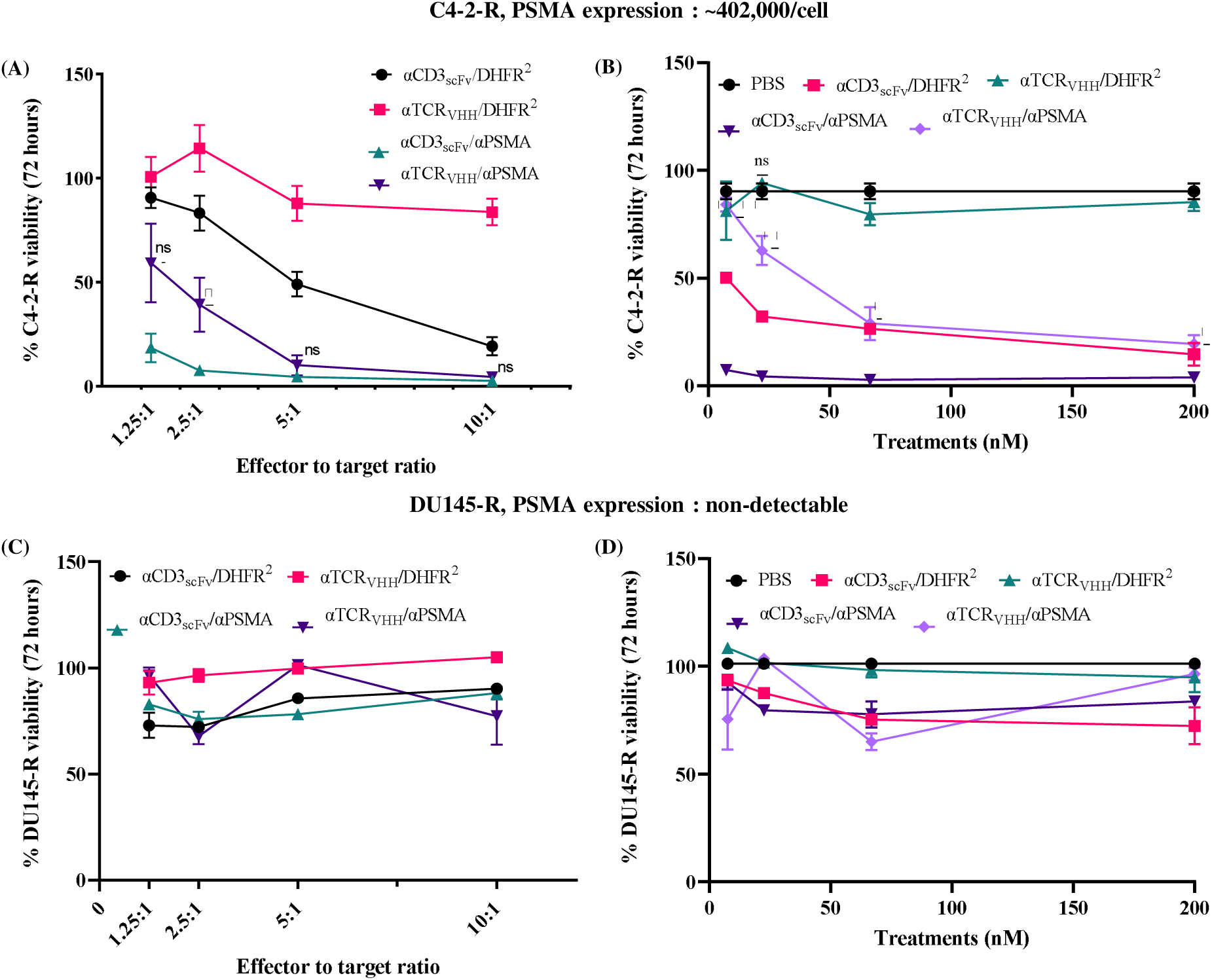
αTCR_VHH_/αPSMA Bispecific CSANs direct cytotoxicity selectively against PSMA+ cell lines. Tumor cells were seeded in 96-well plate as a monolayer. T-cells isolated from healthy donor PBMCs were activated using CD3/CD28 complex and IL-2 and were added to the wells after 20 hours along with monospecific or bispecific CSANs. Tumor cell viability was monitored over 72 hours. Each bullet point represents the final tumor cell count at the end of 72 hour cytotoxicity assay. C4-2-R cell viability (A), and DU145-R cell viability (C) were monitored over 72 hours at a fixed CSAN concentration (100nM) and different E:T ratio. End point cell viability data at each E:T ratio normalized with respect to treatment with PBS only is shown. Significance of αTCR_VHH_/αPSMA was calculated with respect to of αCD3_scFv_/αPSMA by two tailed unpaired t-test. Effects of different concentrations of CSANs on the viability of C4-2-R cells (B), and DU145-R cells (D) were analyzed at a 5:1 E:T ratio. All data is normalized to tumor cells not treated with T-cells or CSANs. Significance of αTCR_VHH_/αPSMA was calculated with respect to of αCD3_scFv_/αPSMA by 2-tailed Student’s *t* test. (Data is displayed as mean ± SD, *P<0.03, **P<0.009, ****P<0.0001). Data shown here is from one donor but is representative of two donors (See Figure S11 in S.I).

### 2.8. Effect of αTCR_VHH_/αPSMA CSANs on T-cell activation

To elucidate the mechanism underlying the observed cytotoxicity towards PSMA^POSITIVE^ C4-2-R cells, we profiled the expression of activation markers on the CD4+ and CD8+ T-cells from the co-culture following treatments with the bispecific constructs and the monospecific controls. While the initial kinetic profiling was attempted at 24 hours, the analysis was optimized for the 48 hour time point given the observed level of T-cell proliferation.

The high affinity αCD3_scFv_/αPSMA CSANs and monospecific αCD3_scFv_CSANs induced substantial upregulation of both CD69 and CD25 on CD4+ (Figure 10A, 10B, Figure S12A and S12B) and CD8+ T-cells (Figure 10D, 10E, Figure S12D and S12E), even in the absence of tumor cells. A similar observation was made for EGFR targeted αCD3_scFv_ CSANs suggesting that CD3 engagement by the CSANs is responsible for the non-specific T-cell activation. In contrast, the αTCR_VHH_/αPSMA CSANs and monospecific αTCR_VHH_ CSANs induced by and large less upregulation of CD69 or CD25 on the T-cells than the αCD3_scFv_ CSANs reflecting the differences in the kinetics of T-cell activation by the two platforms. T-cells stimulated by the αTCR_VHH_/αPSMA CSANs exhibited higher expression of the early activation marker CD69 and a lower expression of the late activation marker CD25 compared to the αCD3_scFv_ /αPSMA CSANs consistent with slower activation kinetics for the αTCR_VHH_/αPSMA CSANs. This is reflected in the lower E:T ratio needed to achieve maximum tumor cell killing with αCD3_scFv_/αPSMA CSANs than αTCR_VHH_/αPSMA CSANs (Figure 10A). Nevertheless, similar levels of cytotoxicity were observed between the αCD3_scFv_/αPSMA CSANs and αTCR_VHH_/αPSMA CSANs, but with significantly less non-specific tumor toxicity observed for the αTCR_VHH_/αPSMA CSANs (Figure 10B). In addition, similar to the EGFR-targeted CSANs, the αTCR_VHH_/αPSMA CSANs induced lower levels of PD-1 upregulation by both CD4+ (Figure 10C and Figure S12C) and CD8+ T-cells (Figure 10F and Figure S12F), consistent with the hyperstimulation associated with strong CD3 binding TCEs.

**Figure 10.**
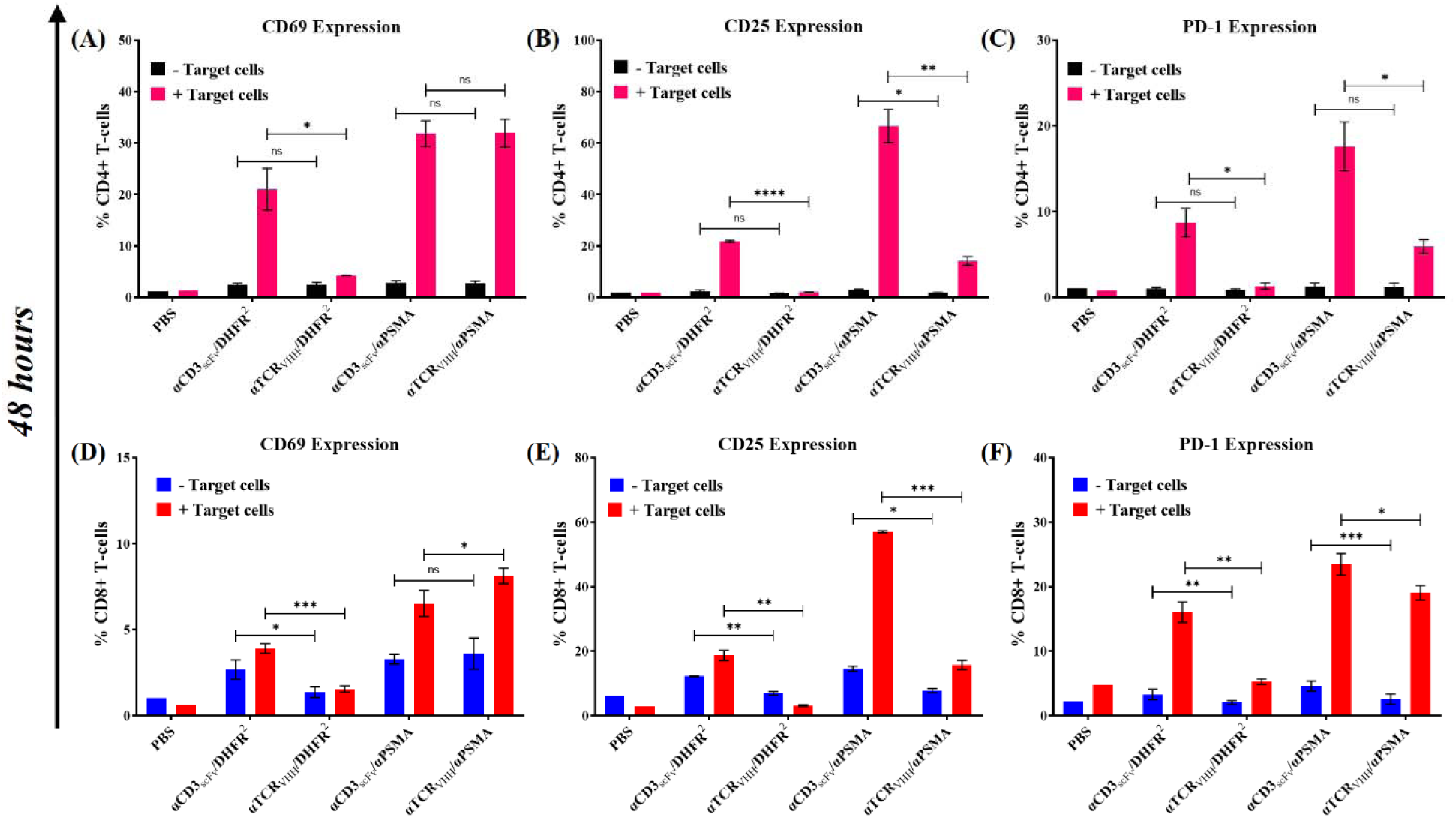
αTCR_VHH_/αPSMA Bispecific CSANs activate T-cells selectively in cytotoxicity assay with C4-2-R cells at 10:1 E:T ratio with 100nM of each treatment. T-cells activated with CD3/CD28 complex and IL-2 were co-cultured with media, monospecific or bispecific CSANs in presence or absence of C4-2-R cells. CD69 expression on CD4+ and CD8+ T cells were measured at 48 hours (A,D). CD25 expression on CD4+ and CD8+ T-cells were measured at 48 hours (B,E) and at. PD-1 expression on CD4+ and CD8+ T-cells were measured at 48 hours (C,F). Data shown is obtained from one donor but is representative of two donors (See Figure S12 in S.I.). Significance for different treatments is calculated by 2-tailed Student’s *t* test. (Data is displayed as mean ± SD, *P<0.05, **P<0.008, ***P<0.0007).

## 3. CONCLUSION

T-cell engagers (TCEs) design has been dominated by the incorporation of a CD3 binder, typically an scFv, since CD3 engagement can initiate activation of the T-cell without the need for co-stimulation. The clinical success of TCEs has been historically associated with two opposing challenges: the need for potent cytotoxicity and the imperative to minimize severe adverse events such as CRS and on-target/ off-tumor toxicity. Furthermore, in the context of solid tumors, the immunosuppressive TME dominated by Regulatory T-cells (Tregs) and the physical barriers to tissue penetration have limited therapeutic efficacy. Previous studies have shown that these issues can be partially addressed by lowering the affinity of the CD3 binder. In this study, we demonstrate that these hurdles could be addressed by shifting the TCE design from high affinity CD3 targeting to lower affinity TCR targeting.

A prevailing belief in immunotherapy has been that higher binding affinity to immune cells is required for superior potency. While this is true, the high reactivity is often accompanied by reduced specificity. We have shown that while the high-affinity αCD3_scFv_ based CSANs (Bispecific as well as monospecific controls) exhibited potent cytotoxicity even at lower dosages, they also displayed “tonic signaling”-activating T-cells and releasing inflammatory cytokines even in the absence of engagement with tumor cell antigen engagement. In contrast, the αTCR_VHH_ CSANs operated under the restriction of “strict engagement.” The moderate affinity of the αTCR_VHH_ was insufficient to trigger T-cell activation autonomously but induced robust lysis upon multivalent cross-linking with tumor cells via the antigens, EGFR and PSMA. This finding suggests that an optimal, intermediate affinity window exists where the T-cell activation signals are not strong enough to push the T-cells to a non-functional exhausted state or weak enough to render them inert, thereby preserving their cytotoxicity, while widening the potential safety margin.^64^

High-affinity engagement with the αCD3_scFv_ CSANs is a known driver of T_reg_ expansion and activation, which can inadvertently shield the tumor from immune attack and contribute to the immunosuppressive microenvironment. Here, we demonstrated that the differential impact of the signaling strength may potentially impact T_reg_ expansion. Lower affinity αTCR_VHH_ CSANs have a potential to promote ‘functional uncoupling’-providing sufficient stimulation for effector T-cell cytotoxicity but falling below the signal strength threshold required for robust T_reg_ expansion. By evading this negative feedback loop, the αTCR_VHH_ based TCEs may sustain anti-tumor immunity within the TME longer than αCD3_scFv_ based TCEs binders. Interestingly, our data shows that this functional uncoupling is not absolute, but rather highly dependent on the heterogeneity of T-cell population in donor T-cells. While one donor showed significant reduction in the T_reg_ population with the αTCR_VHH_ CSANs, no such significant difference was observed for a second donor. This suggests that while signaling strength pay potentially impact T_reg_ expansion, intrinsic heterogeneity in the immune profiles of different donors strongly impact this outcome. In addition, although the CD3 targeted CSAN TCE had greater potency under some conditions than the TCR targeted CSAN TCE in 2D cell culture, the lower affinity αTCR_VHH_ CSAN TCEs were able to induce greater T-cell penetration and cytotoxicity toward 3D spheroids than the higher affinity αCD3_scFv_ CSAN TCEs. These results highlight the importance of evaluating not just targeted tumor cell cytotoxicity but also tumor cell organization on TCE potency. Taken together, our results suggest that TCE designed to target the TCR may offer a viable alternative to CD3 targeting with improved safety and efficacy.

## MATERIALS AND METHODS

### Cell Lines and Cell Culture Conditions

A431, MDA-MB-231, MCF7 and DU145 cells were purchased from the American Type Cell Culture Collection (ATCC). C4-2 cell line was kindly supplied by the lab of Dr. Nicholas Zorko (University of Minnesota). Cell lines were transduced with IncuCyte NucLight green or red Lentivirus Reagents (Sartorius, 4624, and 4625, respectively) following the manufacturers protocol to express the nuclear restricted GFP or mKate2 red fluorescent protein for live-cell imaging assays. The resulting red and green fluorescence labeled cells were named with the suffix ‘R’ and ‘G’ respectively for the purpose of the paper (A431-R, MDA-MB-231-G, MCF7-R, C4-2-R and DU145-R). A431-R, MDA-MB-231-G, MCF7-R, and DU145-R cell lines were cultured in Dulbecco’s modified Eagle’s Medium (DMEM) supplemented with 10% FBS and 1% Glutamax Supplement (Gibco, Catalog number 35050061), at 37°C and 5% CO2. C4-2-R cells were cultured in a Roswell Park Memorial Institute (RPMI) medium supplemented with 10% FBS and 1% Glutamax Supplement (Gibco, Catalog number 35050061), at 37°C and 5% CO2. Human Peripheral Blood Mononuclear Cells (PBMCs) were isolated and purified from the Buffy Coats of blood samples from healthy donors using Lymphocyte Separation Buffer (Corning, Catalog Number: 25-072-CV). Healthy donor blood samples were purchased from Memorial Blood Centers in St. Paul, MN, USA. T-cells were isolated from the PBMCs using the Human T-cell Isolation kit (Akadeum Life Sciences, Catalog Number 13210-120) following the manufacturer’s protocols. The purified T-cells were further activated using ImmunoCult Human CD3/CD28 T-cell Activator Complex (Catalog Number: 10990) and expanded and cultured using ImmunoCult-XF T-Cell Expansion Medium (Stemcell Technologies, Catalog Number: 10981) supplemented with 100 IU/mL IL-2 and 75 μg/mL Normocin following manufacturer’s protocols. The T-cells were suspended in CryoStor®CS10 (Stemcell Technologies, Catalog Number: 210102) media prior to storage.

### Expression Plasmids

The gBlock encoding the gene fragment for αTCR_VHH_-DHFR^2^ fusion protein was ordered form Integrated DNA Technologies (ITD) and cloned into the pET28a vector (EMD Millipore, catalog number: 69864–3) via the NcoI and BamHI restriction sites. The fusion protein was designed with a FLAG and His tag to enable flow cytometric detection for various applications.

### Protein Expression and Purification

The αTCR_VHH_-DHFR^2^ fusion protein was expressed in SHuffle T7 Express Competent *E. coli* cells (New England Biolabs, Catalog Number: C3029J). The *E. coli* cells were cultured in Luria-Bertani (LB) media having 50 μg/mL Kanamycin at 30°C until the OD_600_ reached 0.4-0.6 following which the protein expression was induced using 0.5 mM IPTG at 30°C for 4 hours. The protein was expressed as a soluble protein and purified from soluble fractions of the cell lysate via Immobilized Metal Affinity Chromatography (IMAC) using HisPur™ Cobalt Resin (Thermo Fisher Scientific, Catalog Number: 89965). 150 mM Imidazole was used to elute protein from the resin. The purified protein was then desalted using Amicon® Ultra Centrifugal Filter, 30 kDa MWCO (Milipore, Catalog Number: UFC903024) and analyzed by gel electrophoresis using NuPAGE Bis-Tris protein gels (Thermo Fisher Scientific, Catalog Number: NPO323BOX).

### CSAN formation and Characterization

CSANs were formed by the addition of 2.5-fold molar excess of the chemical dimerizer bisMTX to a solution of DHFR^2^ fusion proteins in PBS (1:1 ratio). The solution was allowed to incubate in dark at room temperature for 1 hour. The monomeric proteins and the CSANs were characterized by analytical Size Exclusion Chromatography on a Superdex G200 column equipped with diode array detector with PBS as the mobile phase to detect the change in the hydrodynamic radius of the CSANs compared to the monomeric proteins. The hydrodynamic radius and polydispersity index of the monomers and CSANs were measured using Dynamic Light Electron Scattering performed on Anton Paar Litesizer 500 (Ashland, VA). 200 μl of 8 μM samples of both monomeric proteins and CSANs were made and filtered using 0.02 μm filter (Cytiva Whatman, Catalog Number: 6809-1002) for measurement. Hydrodynamic radius and polydispersity values are reported as mean ± SD of three measurements.

### Apparent affinity of **α**TCR_VHH_-DHFR^2^ on Jurkat T-cells

The apparent affinity of αTCR_VHH_-DHFR^2^ and αTCR_VHH_ /DHFR^2^ CSANs was determined by flow cytometry. Jurkat T-cells were harvested and washed with PBS buffer with 2% Bovine Serum Albumin (2% BSA/PBS), and 300,000 cell aliquots were resuspended in DPBS buffer containing αTCR_VHH_-DHFR^2^ monomers or αTCR_VHH_ /DHFR^2^ CSANs for 2 hours on ice. The cells were then pelleted and washed with 2% BSA/PBS buffer before incubating with PE-FLAG antibody (BioLegend, Catalog Number: 637309) for 1 hour in dark on ice. After incubation, the cells were washed and resuspended in 200 μL 2% BSA/PBS and analyzed using LSR II Flow Cytometer (BD Biosciences)

### *In vitro* cytotoxicity Assays

Loss of GFP or mKate2 fluorescence was monitored using BioTek Cytation C10 Confocal Imaging Reader (Agilent) equipped with BioTek BioSpa 8 Automated Incubator to report real time cancer cell death. One day prior to the cytotoxicity assay, 5 x 10^3^ A431-R cells, 10 x 10^3^ MDA-MB-231-G, or 10 x 10^3^ MCF7-R cells were seeded into 96-well plates in 100 μL of DMEM media per well. A431-R spheroids were formed by seeding 2500 cells/well in Nunclon™ Sphera™ 96-well Ultra Low Attachment U-Bottom Plate followed by centrifugation at 150xg for 5 min 24 hours prior to the experiment. Previously purified human PBMCs were thawed and allowed to rest in RPMI supplied with 75 μg/mL Normocin overnight at 37°C and with 5% CO_2_. 18-20 hours later, T-cells were isolated from the PBMCs following manufacturer’s protocol and counted and resuspended in RPMI. Mono-specific and Bispecific CSANs were freshly made in PBS (25 μL per well of the indicated concentration) and incubated with the appropriate number of freshly isolated T-cells in RPMI (determined by E:T ratio, 75 μL per well) for 1 hour at 37°C with 5% CO_2._ Following this incubation, 100 μL of this T-cell and CSAN suspension were added to each well. The plate was placed in BioTek BioSpa 8 Automated Incubator at 37°C and 5% CO_2_ and mKate2 or GFP florescence was measured every 4 hours for 72 hours to determine viability of cancer cells. Cytotoxicity assay on A431-R spheroids was performed similarly. 5 x 10^3^ A431-R cells were seeded in Nunclon™ Sphera™ 96-well Ultra Low Attachment U-Bottom Plate (Catalog Number: 174925) and centrifuged at 150xg for 5 minutes to form the spheroids 18-20 hours prior to treatment. For the PSMA positive cell lines, 10 x 10^3^ C4-2-R cells and 5 x 10^3^ DU145-R cells were seeded in 96-well plates in 100 μL of RPMI and DMEM respectively, 1 day prior to the assay. Previously isolated and expanded T-cells were thawed and resuspended in RPMI with 75 μg/mL Normocin and rested overnight in 37°C with 5% CO_2._ The following day, the T-cells were washed and counted and T-cells-CSAN suspension was prepared as mentioned above. All images were obtained at 4X objective. Each cytotoxicity graph was obtained with n=3 wells per time point using T-cells from one donor but is representative of two different donors. The cell viability at each time point is shown as mean ± SD. For A431-R spheroids, images were obtained at 20X objective and z-projection images are reported.

### Cytokine Analysis by ELISA

The supernatants from the target cell – T-cell coculture plates were collected at 24 hours and 72 hours from two separate plates. IFN-γ and TNF-α concentrations were quantified using Sandwich ELISA according to manufacturer’s protocol (BD OptEIA Human IFN-γ ELISA set, Catalog Number: 555142, and BD OptEIA Human TNF ELISA set, Catalog Number: 555212)

### T-cell activation analysis using flow cytometry

After 24 hours and 72 hours, 96-well plates from the target cell-T cell co-culture studies were centrifuged at 400xg for 5 minutes, supernatants were discarded and the remaining T-cells were resuspended in 100 μL RPMI per well. T-cell activation markers were stained with anti-human CD4 (BioLegend, Cat Number: 334606, Clone SK3), anti-human CD8 (BioLegend, Cat Number: 344756, Clone SK1), anti-human CD25 (BioLegend, Cat Number: 356106, Clone M-A251), anti-human CD69 (BioLegend, Cat Number: 985206, Clone FN50), anti-human PD-1 (BioLegend, Cat Number: 329919, Clone EH12.2H7) and anti-human CD127 (BioLegend, Cat Number: 351310, Clone A019D5). Memory T-cell analysis was performed by staining with anti-human CD4, anti-human CD8, anti-human CD45RO (BioLegend, Cat Number: 304224, Clone UCHL1) and anti-human CCR7 (BioLegend, Cat Number: 353216, Clone G043H7). Staining solution was made by mixing the manufacture recommended volume of each antibody or its isotype in PBS containing 2% BSA/PBS. 50 μL of the staining solution was added to each well in the 96-well plate and incubated in dark for 1 hour on ice. All flow cytometry analysis was done using Beckman Coulter CytoFLEX S Flow Cytometer and values are displayed as the mean ± SD of at least 3 replicates from one donor but is representative of two donors.

## Author Contributions

C.R.W and D.P conceived and designed the project. D.P designed and generated the αTCR_VHH_-DHFR^2^ protein, performed all the experiments and analyses presented in the paper and wrote the manuscript. αPSMA and αEGFR (E1) were generated by A.K and D.G.D respectively. BisMTX was synthesized by F.R. L.R prepared the RFP and GFP cancer cell lines. All authors provided edits and comments.

## ACKNOWLEDGMENT

We gratefully acknowledge support from NCI R01CA247681 (CRW), the University of Minnesota Cancer Center and University of Minnesota Foundation.

## ABBREVIATIONS

APC: antigen-presenting cells
ATCC: American Type Culture Collection
B-ALL: B-cell precursor acute lymphoblastic leukemia
BSA: bovine serum albumin
CDR3: third complementarity-determining region
CRS: cytokine release syndrome
CSAN: chemically self-assembled nanoring
DCs: dendritic cells
DLS: dynamic light scattering
DMEM: Dulbecco’s modified Eagle medium
DPBS: Dulbecco’s phosphate-buffered saline
EGFR: epidermal growth factor receptor
ELISA: enzyme-linked immunosorbent assay
E:T ratio: effector-to-target ratio
FBS: fetal bovine serum
FLAG: FLAG epitope tag (DYKDDDDK)
FR: framework region
GFP: green fluorescent protein
IFN: interferon
IL: interleukin
IMAC: immobilized metal affinity chromatography
IPTG: isopropyl β-D-1-thiogalactopyranoside
ISV: immunoglobulin single variable domain
LB: Luria–Bertani
MDSCs: myeloid-derived suppressor cells
MHC: major histocompatibility complex
MWCO: molecular weight cutoff
mKate2: monomeric far-red fluorescent protein Kate2
NK: natural killer
PBS: phosphate-buffered saline
PBMC: peripheral blood mononuclear cell
PD-1: programmed cell death protein 1
PE: phycoerythrin
PSMA: prostate-specific membrane antigen
RPMI: Roswell Park Memorial Institute medium
scFv: single-chain variable fragment
SD: standard deviation
SEC: size exclusion chromatography
TAA: tumor-associated antigen
TCE: T-cell engager
TCR: T-cell receptor
TGF: transforming growth factor
TME: tumor microenvironment
TNF: tumor necrosis factor
VHH: variable domain of heavy-chain-only antibody.

