## Supplemental File for "T-cell Receptor (TCR) Targeting with Multivalent T-cell Engagers"

2231 6th Street S.E.

Cancer & Cardiovascular Research Building

Minneapolis, Minnesota 55455, USA

**Table of Contents**


Supplementary Figure S1. Expression and purification of αTCR$_{VHH}$/DHFR$^2$……………….…………….S3

Supplementary Figure S2. Donor #2: αTCR$_{VHH}$/αEGFR (E1) Bispecific CSANs direct cytotoxicity against EGFR+ cell lines……………………………………………………………………………..…………….S4

Supplementary Figure S3. Donor #2: αTCR$_{VHH}$/αEGFR (E1) Bispecific CSANs activate T-cells selectively in cytotoxicity assay with A431-R cells ………………………………………..……………………S5

Supplementary Figure S4. Donor #2: αTCR$_{VHH}$/αEGFR (E1) Bispecific CSANs activate T-cells selectively in cytotoxicity assay with MDA-MB-231-G cells …………………………..…………………………S6

Supplementary Figure S5. Donor #1 and Donor #2: Memory T-cell formation by αTCR$_{VHH}$/αEGFR (E1) Bispecific CSANs in cytotoxicity assay with A431-R cells ………………………………………………S7

Supplementary Figure S6. Donor #1 and Donor #2: Memory T-cell formation by αTCR$_{VHH}$/αEGFR (E1) Bispecific CSANs in cytotoxicity assay with MDA-MB-231-G cells ……………………………………S8

Supplementary Figure S7. Donor #2: Regulatory T-cell response to CSANs *in vitro* ..……………………S9

Supplementary Figure S8. αTCR$_{VHH}$/αEGFR (E1) Bispecific CSANs directed end point MDA-MB-231-G cell viability at 24 hours and 72 hours …………………………………………………………………S10

Supplementary Figure S9. Donor #2: Determination of *in vitro* cytokine release as a result of αTCR$_{VHH}$/αEGFR (E1) mediated MDA-MB-231-G cell lysis……..…………………………………..……S11

Supplementary Figure S10. Donor #2: αTCR$_{VHH}$ /αEGFR (E1) Bispecific CSANs induced cytotoxicity against 3D spheroids of A431-R cells ……………………………….…………………………………………………S12

Supplementary Figure S11. Donor #2: αTCR$_{VHH}$/αPSMA Bispecific CSANs direct cytotoxicity selectively against PSMA+ cell lines……………………………………...………………………………………S13

Supplementary Figure S12. Donor #2: αTCR$_{VHH}$/αPSMA Bispecific CSANs activate T-cells selectively in cytotoxicity assay with C4-2-R cells ……………………………………………….…………………S14

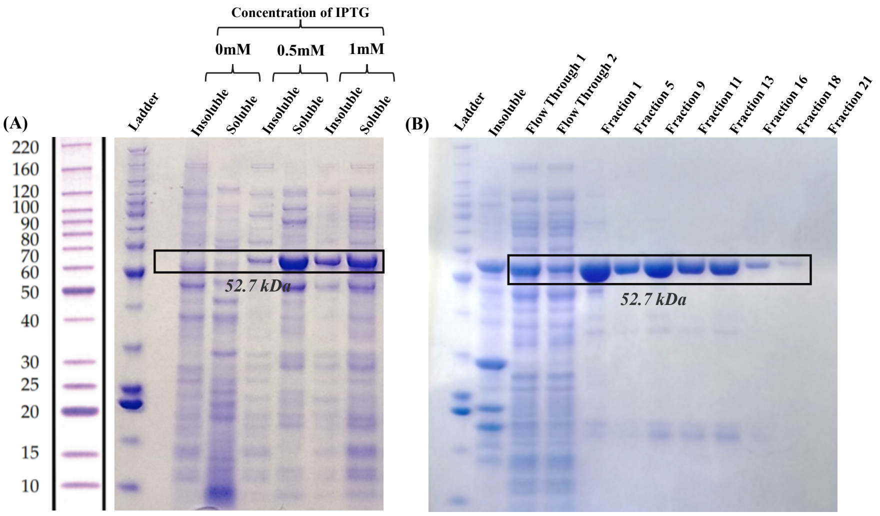


**Figure S1 Expression and purification of αTCR$_{VHH}$/DHFR$^2$. (A)** Expression of αTCR$_{VHH}$/DHFR$^2$ at different concentrations of IPTG. **(B)** SDS-PAGE of αTCR$_{VHH}$/DHFR$^2$ after purification using Cobalt Column.

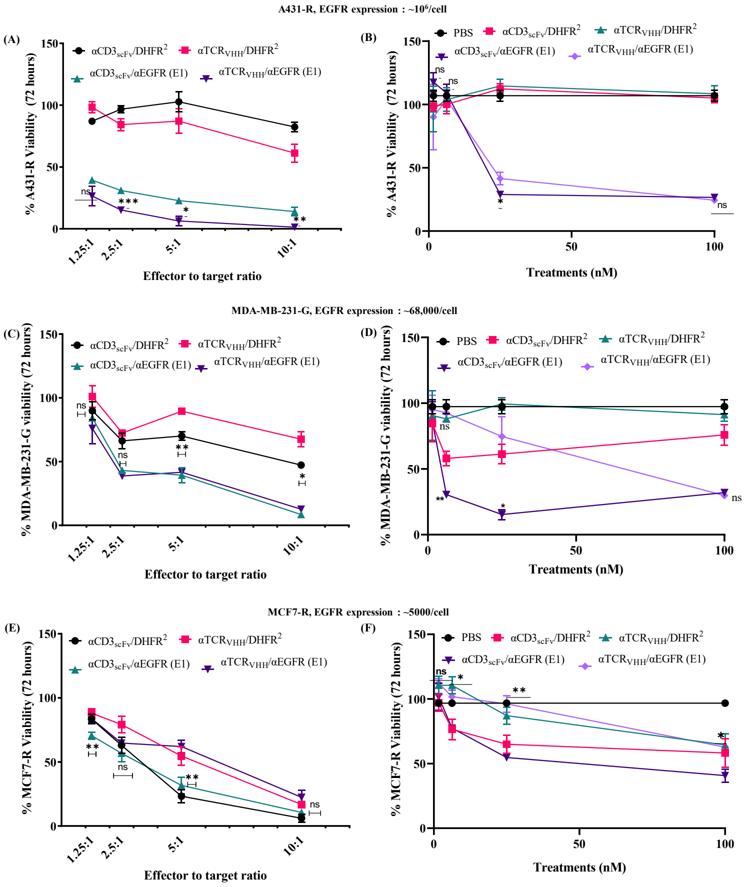


**Figure S1. αTCR$_{VHH}$/αEGFR (E1) Bispecific CSANs direct cytotoxicity against EGFR+ cell lines**. Tumor cells were seeded in 96-well plate as a monolayer. T-cells freshly isolated from healthy donor PBMCs were added to the wells 20 hours after with monospecific or bispecific CSANs. Tumor cell viability was monitored over 72 hours. Each bullet point represents the final tumor cell count at the end of 72 hour cytotoxicity assay. A431-R cell viability **(A)**, MDA-MB-231-G cell viability **(C)**, and MCF7-R cell viability **(E)** were monitored over 72 hours at a fixed CSAN concentration (100nM) and different E:T ratio. End point cell viability data at each E:T ratio normalized with respect to treatment with PBS only is shown. Significance of αTCR$_{VHH}$/αEGFR (E1) was calculated with respect to αCD3scFv/αEGFR (E1) by two tailed unpaired t-test **(A)** and **(E)**. Significance of αTCR$_{VHH}$/DHFR$^2$ was calculated with respect to αCD3scFv/ DHFR$^2$ **(C)** by two tailed unpaired t-test. Effects of different concentrations of CSANs on the viability of A431-R cells **(B)**, MDA-MB-231-G cells **(D)** and MCF7-R cells **(F)** were analyzed at a 5:1 E:T ratio. All data is normalized to tumor cells not treated with T-cells or CSANs. Significance of αTCR-DHFR$^2$/E1-DHFR$^2$ was calculated with respect to αCD3scFv/ αEGFR (E1) by two tailed unpaired t-test. Data shown is obtained from one donor but is representative of two donors (Figure **2** and **S2**).

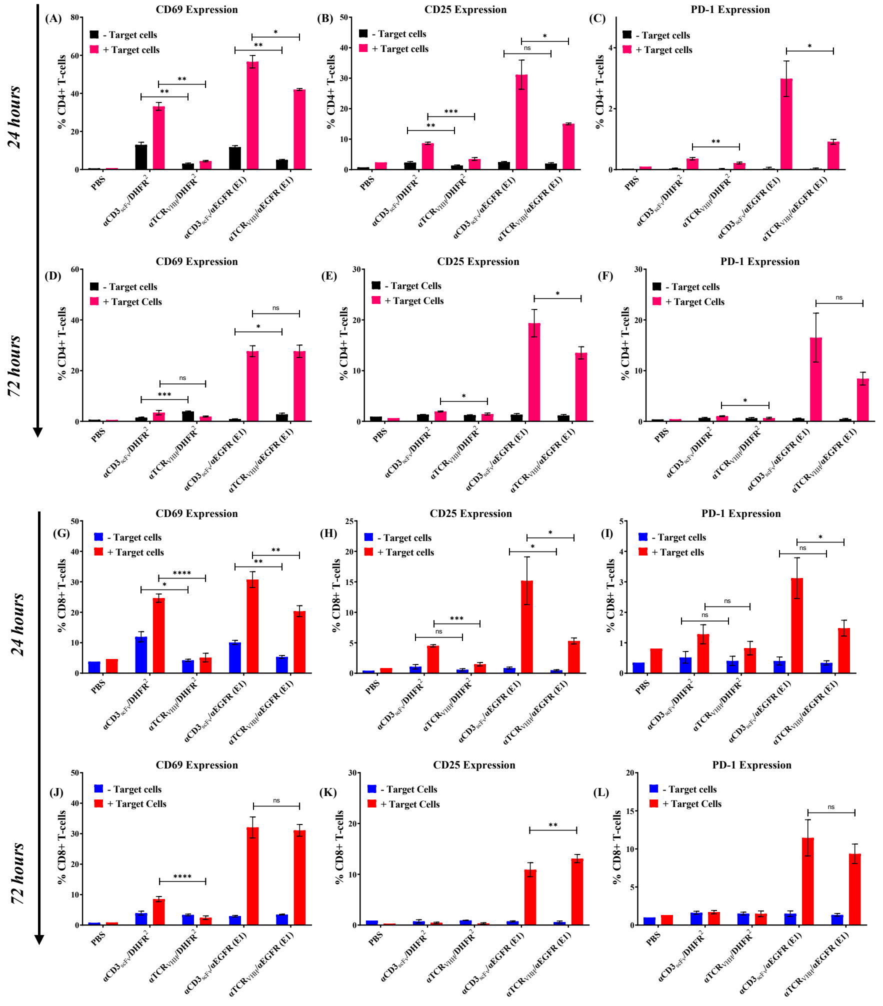


**Figure S3. αTCR$_{VHH}$/αEGFR (E1) Bispecific CSANs activate T-cells selectively in cytotoxicity assay with A431-R cells**. A431-R cells were seeded into half of a 96-well plate as a monolayer. 20 hours later, T-cells freshly isolated from healthy donor PBMCs were co-cultured with media, monospecific or bispecific CSANs (100nM) at 10:1 E:T ratio in presence or absence of A431-R cells. CD69 expression on CD4+ and CD8+ T cells were measured at 24 hours **(A,G)** and at 72 hours **(D,J)**. CD25 expression on CD4+ and CD8+ T-cells were measured at 24 hours **(B,H)** and at 72 hours **(E,K)**. PD-1 expression on CD4+ and CD8+ T-cells were measured at 24 hours **(C,I)** and at 72 hours **(F,L)**. Data shown is obtained from one donor but is representative of two donors (Figure **3** and **S3**). Significance for different treatments is calculated by 2-tailed Student's t test.

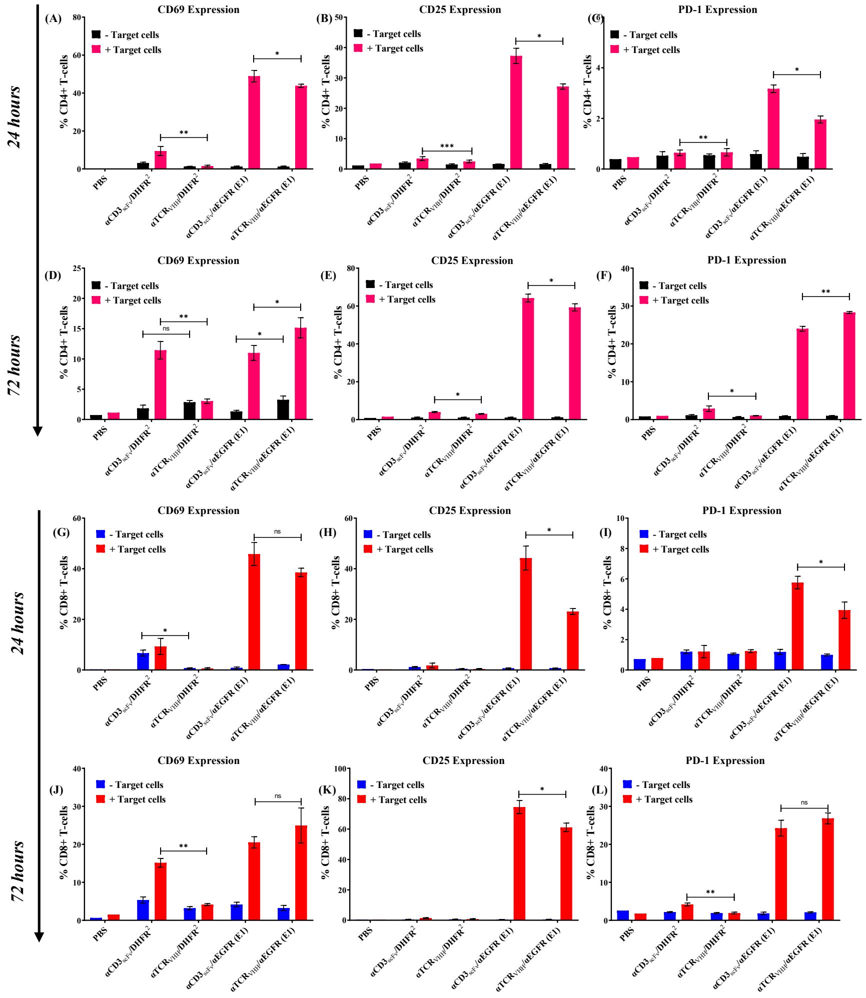

**Figure S4. αTCR$_{VHH}$/αEGFR (E1) Bispecific CSANs activate T-cells selectively in cytotoxicity assay with MDA-MB-231-G cells**. MDA-MB-231-G cells were seeded into half of a 96-well plate as a monolayer. 20 hours later, T-cells freshly isolated from healthy donor PBMCs were co-cultured with media, monospecific or bispecific CSANs (100nM) at 5:1 E:T ratio in presence or absence of MDA-MB-231-G cells. CD69 expression on CD4+ and CD8+ T cells were measured at 24 hours **(A,G)** and at 72 hours **(D,J)**. CD25 expression on CD4+ and CD8+ T-cells were measured at 24 hours **(B,H)** and at 72 hours **(E,K)**. PD-1 expression on CD4+ and CD8+ T-cells were measured at 24 hours **(C,I)** and at 72 hours **(F,L)**. Data shown is obtained from one donor but is representative of two donors (Figure **4** and **S4**). Significance for different treatments is calculated by 2-tailed Student's t test.

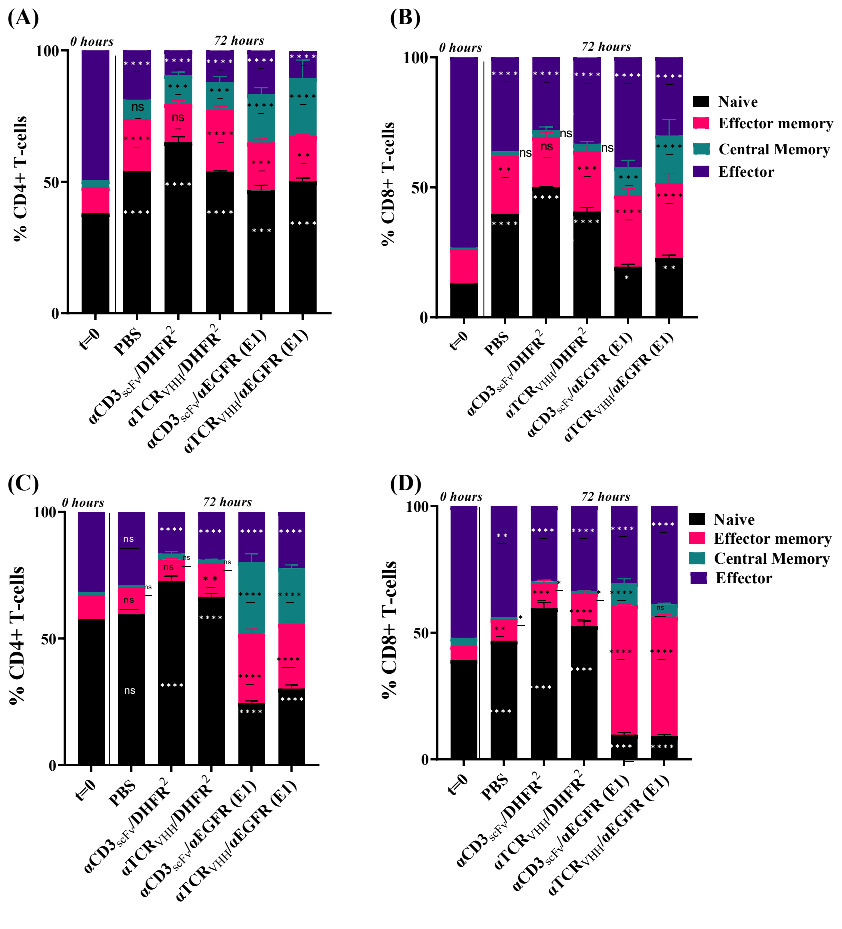


**Figure S5. Memory T-cell formation by $\alpha$TCR$_{VHH}$/$\alpha$EGFR (E1) Bispecific CSANs in cytotoxicity assay with A431-R cells**. Changes in the population of CD4+ and CD8+ naïve, memory and effector T-cells activated by Monospecific and Bispecific CSANs in cytotoxicity assay with A431-R cells at 10:1 E:T ratio with 100nM of each treatment were measured using flow cytometry. T-cells isolated from healthy donor PBMCs were co-cultured with media, monospecific or bispecific CSANs in presence of MDA-MB-231-G cells. CD4+ and CD8+ T-cell populations from donor 1 **(A,B)** and donor 2 **(C,D)** were characterized after 72 hours of co-culture assay by measuring the expression of CD45RO and CCR7 using flow cytometry. The T-cell phenotypes are described as: naïve (CD45RO$^-$/CCR7$^+$), central memory (CD45RO$^+$/CCR7$^+$), effector memory (CD45RO$^+$/CCR7$^-$) and effector (CD45RO$^-$/CCR7$^+$). Significance for different treatments is calculated with respect to the population of freshly isolated T-cells at 0 hours by 2-way Anova.

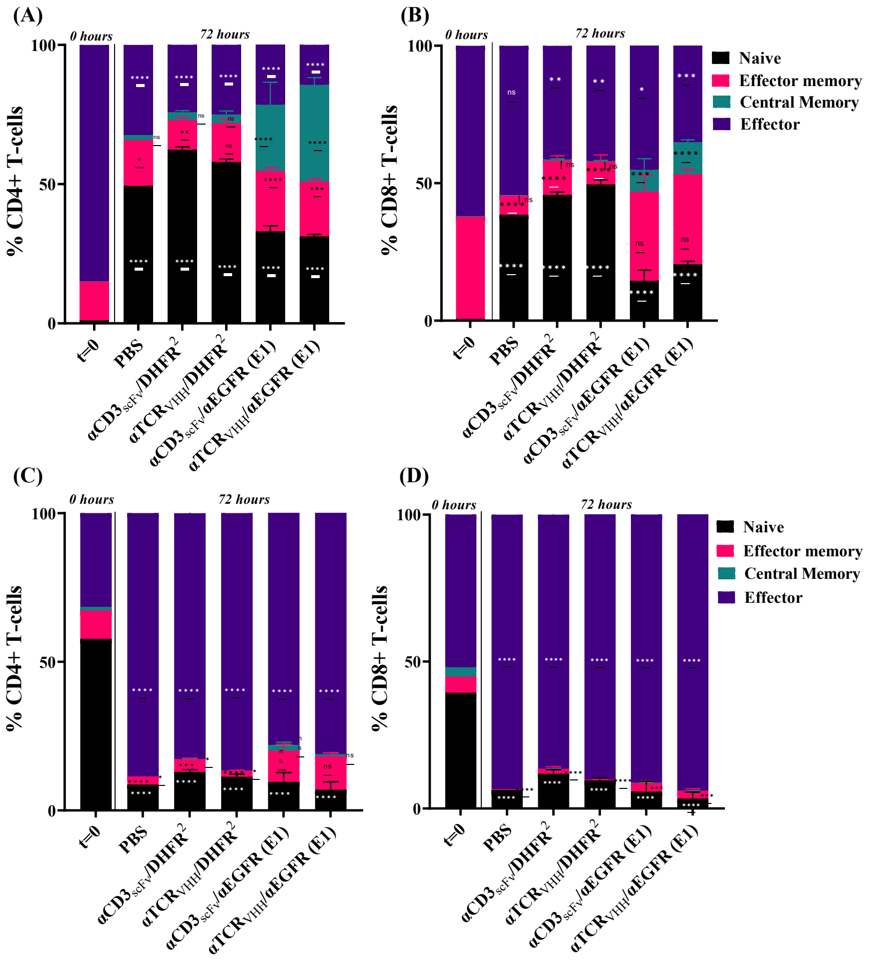


**Figure S6. Memory T-cell formation by αTCR$_{VHH}$/αEGFR (E1) Bispecific CSANs in cytotoxicity assay with MDA-MB-231-G cells**. Changes in the population of CD4+ and CD8+ naïve, memory and effector T-cells activated by Monospecific and Bispecific CSANs in cytotoxicity assay with MDA-MB-231-G cells at 5:1 E:T ratio with 100nM of each treatment were measured using flow cytometry. T-cells isolated from healthy donor PBMCs were co-cultured with media, monospecific or bispecific CSANs in presence of MDA-MB-231-G cells. CD4+ and CD8+ T-cell populations from donor 1 **(A,B)** and donor 2 **(C,D)** were characterized after 72 hours of co-culture assay by measuring the expression of CD45RO and CCR7 using flow cytometry. The T-cell phenotypes are described as: naïve (CD45RO$^-$/CCR7$^+$), central memory (CD45RO$^+$/CCR7$^+$), effector memory (CD45RO$^+$/CCR7$^-$) and effector (CD45RO$^-$/CCR7$^+$). Significance for different treatments is calculated with respect to the population of freshly isolated T-cells at 0 hours by 2-way Anova.

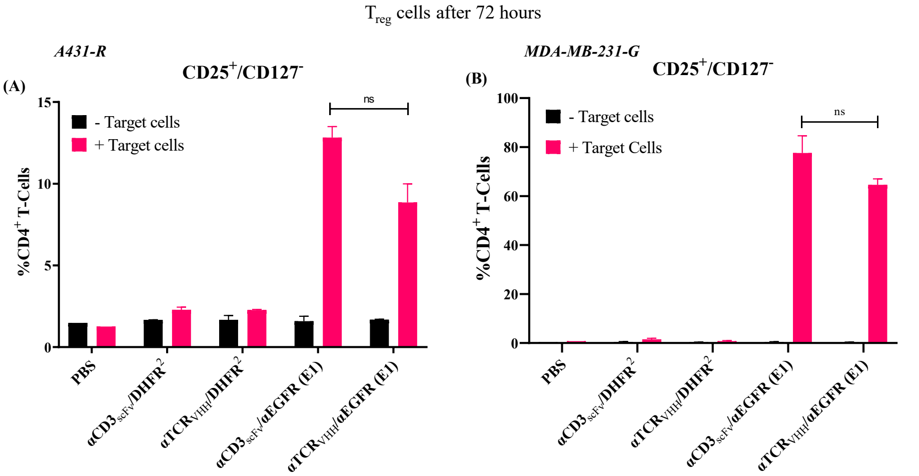


**Figure S7. Regulatory T-cell response to CSANs *in vitro*.** T-cells freshly isolated from healthy donor PBMCs were co-cultured with media, monospecific or bispecific CSANs (100nM) in presence or absence of a 2D monolayer of A431-R **(A)** and MDA-MB-231-G **(B)** cells. Following 72 hour incubation, T-cells were harvested and analyzed for CD4, CD25 and CD127 expression. Data shown is obtained from one donor but is representative of two donors (Figure **5**). Significance was calculated by using 2-tailed Student's t test.

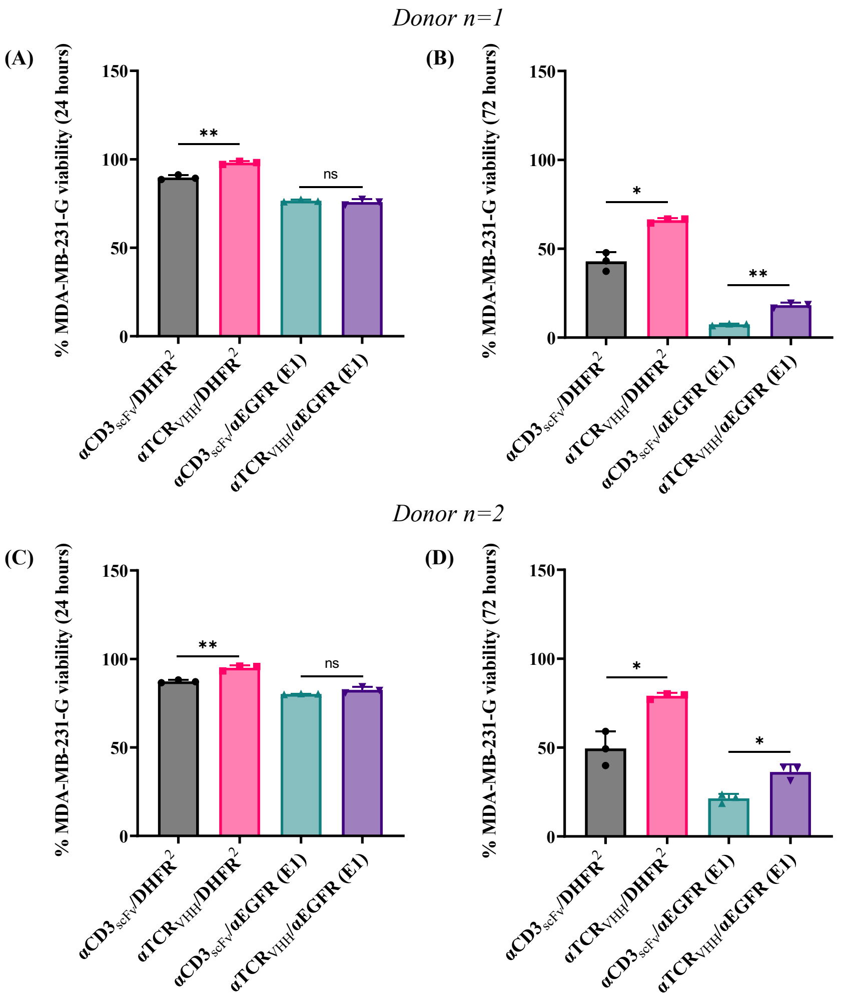


**Figure S8. αTCR$_{VHH}$/αEGFR (E1) Bispecific CSANs directed end point MDA-MB-231-G cell viability at 24 hours and 72 hours.** Tumor cells were seeded in 96-well plate as a monolayer. T-cells freshly isolated from healthy donor PBMCs were added to the wells 20 hours after with monospecific or bispecific CSANs (100nM treatments) at 5:1 E:T ratio. Tumor cell viability was monitored over 24 hours and 72 hours following which supernatants were harvested and used for cytokine analysis. End point cell viability data normalized with respect to treatment with PBS only is shown. Significance was calculated by two tailed unpaired t-test. Data shown is obtained from 2 donors **(A,B)** and **(C,D).**

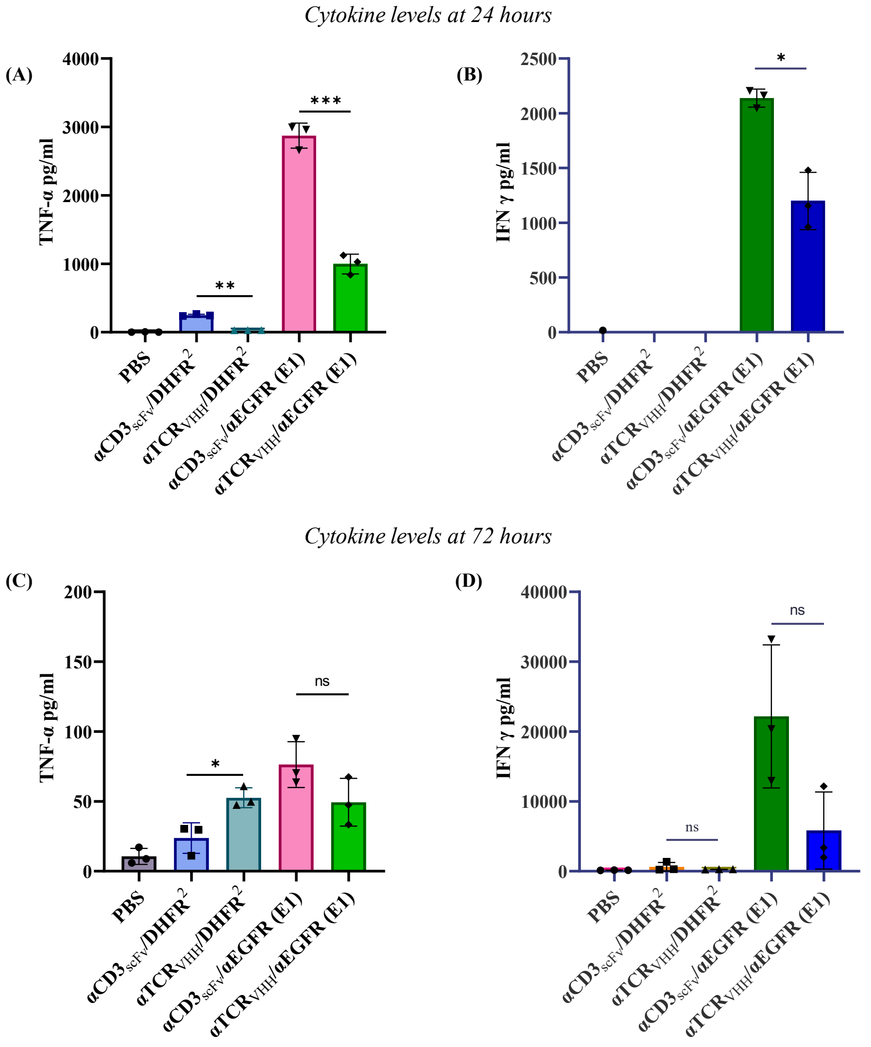


**Figure S9. Determination of *in vitro* cytokine release as a result of αTCR$_{VHH}$/αEGFR (E1) mediated MDA-MB-231-G cell lysis**. Supernatants from MDA-MB-231-G and freshly isolated T-cell co-culture as described in **Figure S8** were analyzed for TNF-α and IFN-γ at 24 hours **(A,B)** and 72 hours **(C,D)** using a sandwich ELISA. Significance was calculated using 2 tailed Student's t test. Data shown is obtained from one donor but is representative of two donors (Figure **6** and **S9**).

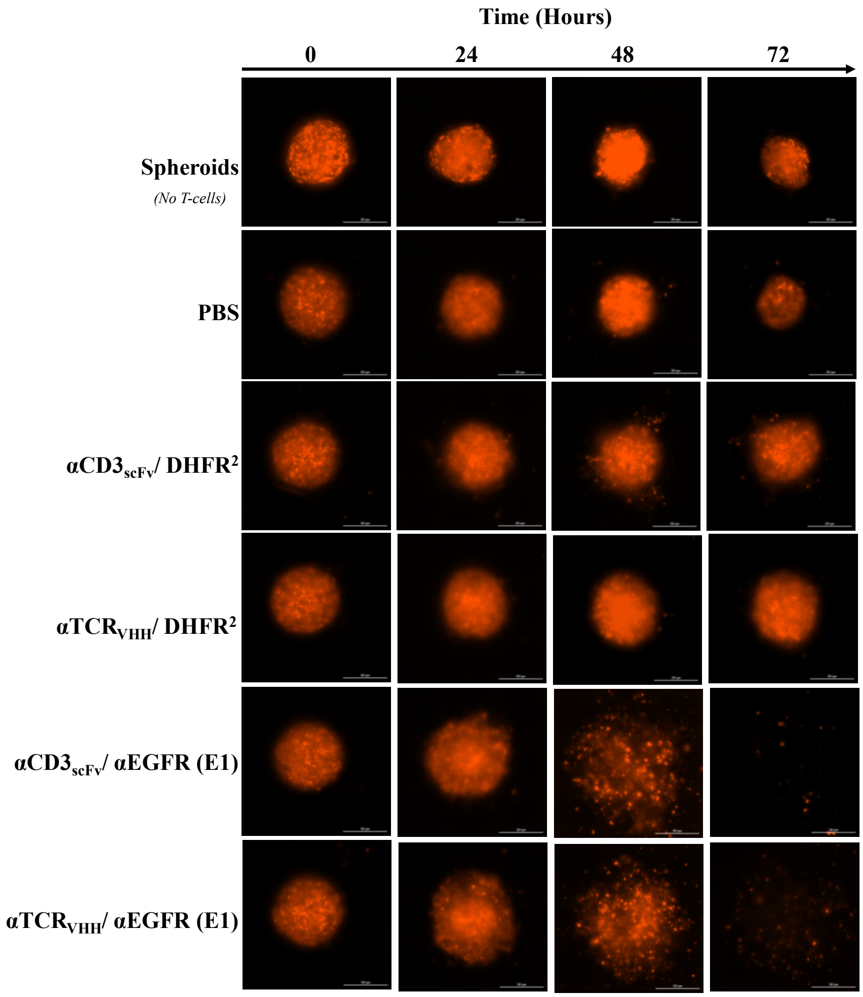


**Figure S10. α$TCR_{VHH}$ /αEGFR (E1) Bispecific CSANs induced cytotoxicity against 3D spheroids of A431-R cells.** Spheroids were incubated with freshly isolated T-cells from healthy donor PBMCs at 10:1 E:T ratio along with PBS, monospecific or bispecific CSANs at 100nM for 72 hours at 37°C and images were taken every 24 hours. Representative confocal images of the spheroids at different times points. Data shown is obtained from one donor but representative of two donors **(Figure 7).**

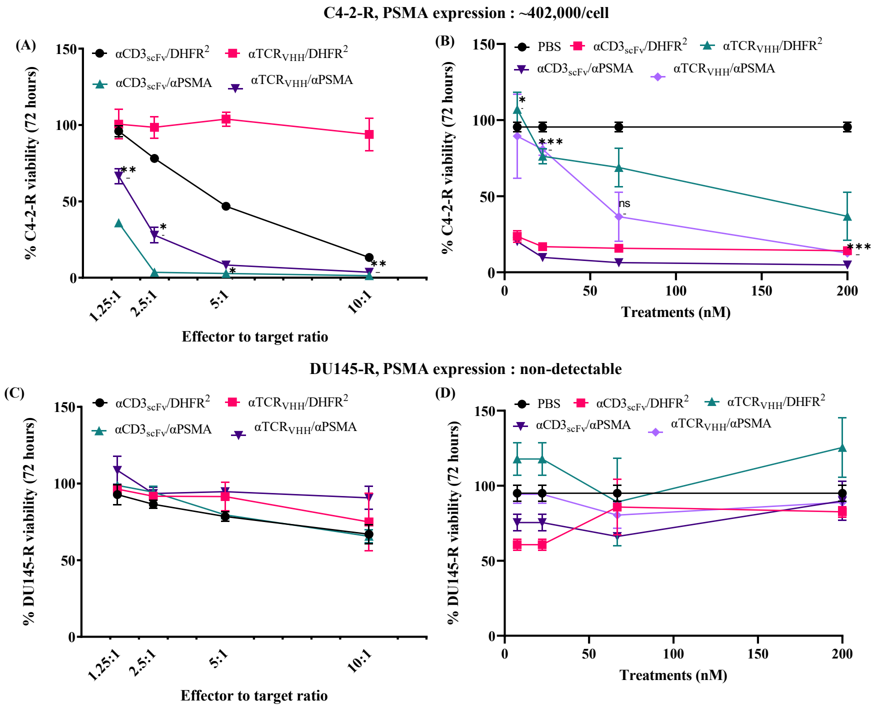


**Figure S11. αTCR$_{VHH}$/αPSMA Bispecific CSANs direct cytotoxicity selectively against PSMA+ cell lines**. Tumor cells were seeded in 96-well plate as a monolayer. T-cells isolated from healthy donor PBMCs were activated using CD3/CD28 complex and IL-2 and were added to the wells 20 hours later with monospecific or bispecific CSANs. Tumor cell viability was monitored over 72 hours. Each bullet point represents the final tumor cell count at the end of 72 hour cytotoxicity assay. C4-2-R cell viability **(A)**, and DU145-R cell viability **(C)** were monitored over 72 hours at a fixed CSAN concentration (100nM) and different E:T ratio. End point cell viability data at each E:T ratio normalized with respect to treatment with PBS only is shown. Significance of αTCR$_{VHH}$/αPSMA was calculated with respect to αCD3$_{scFv}$/αPSMA by two tailed unpaired t-test. Effects of different concentrations of CSANs on the viability of C4-2-R cells **(B)**, and DU145-R cells **(D)** were analyzed at a 5:1 E:T ratio. All data is normalized to tumor cells not treated with T-cells or CSANs. Significance of αTCR$_{VHH}$/αPSMA was calculated with respect to αCD3$_{scFv}$/αPSMA by two tailed unpaired t-test. Data shown is obtained from one donor but is representative of two donors (Figure **8** and **S11**).

**Commented [CW1]:** A has wrong x-axis legend. What is the concentration of CSANs used for A dn C? The legend is confusing.

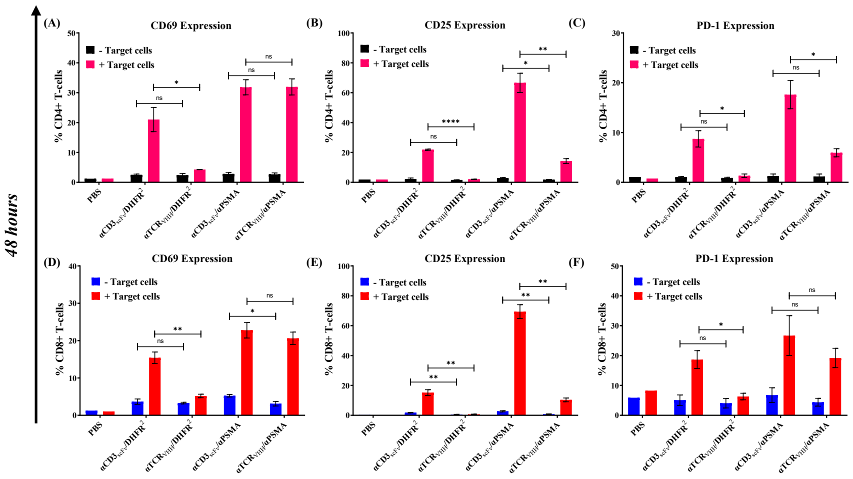


**Figure S12. αTCR$_{VHH}$/αPSMA Bispecific CSANs activate T-cells selectively in cytotoxicity assay with C4-2-R cells**. C4-2-R cells were seeded into half of a 96-well plate as a monolayer. 20 hours later, T-cells activated with CD3/CD28 and IL-2 were co-cultured with media, monospecific or bispecific CSANs (100nM) at 10:1 E:T ratio in presence or absence of C4-2-R cells. CD69 expression on CD4+ and CD8+ T cells were measured at 48 hours **(A,D)**. CD25 expression on CD4+ and CD8+ T-cells were measured at 48 hours **(B,E)** and at. PD-1 expression on CD4+ and CD8+ T-cells were measured at 48 hours **(C,F)**. Data shown is obtained from one donor but is representative of two donors (Figure **9** and **S12**). Significance for different treatments is calculated by 2-tailed Student's t test.
